# Accessible Gibbs energy at metabolic activation limits long-term cell growth

**DOI:** 10.64898/2026.04.30.722108

**Authors:** Yan B. Barreto, Elwin P.H. Jongman, Miyer F. Patiño-Ruiz, Douwe A.J. Grundel, Melike Uysal, Jelmer Coenradij, Bert Poolman, Matthias Heinemann

## Abstract

When exposed to a nutrient, cells activate metabolism by reorganizing metabolite pools and enzyme expression to approach the maximal growth rate permitted by physicochemical constraints. While these constraints define reachable steady states, here we propose that the Gibbs energy accessible at activation further limits which states are reached. Using minimal metabolic models, we find that limited accessible Gibbs energy can trap cells in low-growth states by constraining metabolic reorganization and imposing a proteomic burden on transport and phosphorylation reactions. To investigate this experimentally, we reconstituted the arginine deiminase pathway in vesicles, revealing that the size of a conserved pool of interconverting metabolites (arginine, citrulline, and ornithine) determines accessible Gibbs energy and constrains steady-state ATP production rate, a proxy for growth. Together, these results indicate that cellular metabolism retains memory of its initial energetic state, with accessible Gibbs energy at activation acting as a thermodynamic constraint on long-term growth.

## Introduction

Cells adapt to environmental changes by reorganizing metabolic fluxes and reallocating proteomic resources, transitioning from one non-equilibrium steady state to another.^1–3^ Traditionally, these metabolic states have been assumed to be independent, with emergent properties such as growth rate determined solely by current conditions. This assumption underlies many theoretical descriptions of cellular metabolism, from growth laws such as the Monod equation to constraint-based approaches such as flux balance analysis.^4^ However, recent findings suggest that a cell’s past metabolic states can influence present cellular responses, introducing the notion of metabolic memory.^5–9^ Such memory can arise when transient metabolic states leave behind altered metabolite pools^10^ or impose proteome-level constraints,^11^ which in turn shape subsequent growth behaviors such as lag times and adaptation rates. These findings raise a fundamental question: how does a cell’s metabolic state during transient processes influence its long-term growth capacity?

One such transient process is the activation of metabolism after starvation, where many organisms rely on “turbo” pathways, a metabolic design that requires an upfront investment of energy before energy is gained again. Glycolysis provides a well-known example, consuming two ATP molecules before producing four ATP through glucose catabolism. The arginine deiminase (ADI) pathway also follows a turbo design: it dissipates the chemical gradient established by intracellular ornithine to import arginine, and the imported arginine is then broken down to regenerate ornithine and produce one ATP molecule. The turbo design may offer an evolutionary advantage by enabling a more rapid flux generation once metabolism is activated.^12^ However, it also creates a vulnerability: because activation requires an upfront energetic investment, it can fail if the cell’s initial energetic state cannot support this investment.^13, 14^ While this risk has been recognized, broader implications of the turbo design for cellular growth remain largely unexplored. In particular, it is unclear whether the initial energetic state also influences the growth capacity of cells that successfully resume metabolism. In this paper, we hypothesize that the initial energetic state of a cell leaves a lasting imprint on its long-term growth. Specifically, we propose that the amount of Gibbs energy available in the initial metabolic state constrains the cell’s steady-state growth rate.

To test this hypothesis, we constructed and analyzed a model of a cell activating its metabolism and transitioning to steady growth, incorporating turbo-designed mechanisms commonly used by cells. The model shows that, unlike classical growth models such as the Monod equation,^15^ which link growth to external nutrient levels alone, identical cells initialized with different amounts of Gibbs energy grow at distinct rates despite identical environments. We show that this behavior results from the pathway’s turbo design, which provides the system with memory of its history. To validate the model’s predictions, we reconstituted the ADI pathway, the simplest known turbo-designed system, in vesicles initiated with different Gibbs energy levels. Experiments confirm that ATP production rate, a proxy for growth rate in the experimental system, is governed by the buildup of an intermediate metabolite that links the initial Gibbs energy levels to the vesicles’ long-term metabolic capacity, in agreement with the model predictions. Together, these results show that the Gibbs energy available at metabolic activation constrains long-term cellular growth, indicating that the steady-state behavior of cellular metabolism can retain memory of its initial energetic state.

## Results

### Development of a cell model incorporating the turbo design

To test whether the amount of Gibbs energy available at an initial metabolic state constrains steady-state growth, we extended the coarse-grained cell model of Noor et al.^16^ to include the turbo design. Specifically, our model represents metabolism as a sequence of four reactions (Fig. 1a): import of an external nutrient (denoted as reaction “t”); phosphorylation of the imported nutrient (reaction “p”); catabolism generating ATP and a precursor (reaction “c”); and ATP-driven anabolism producing biomass from the precursor (reaction “a”). In Noor et al.’s model, nutrient transport and phosphorylation are lumped into a single uptake reaction driven by the external nutrient concentration. In contrast, our extension separates transport and phosphorylation and also includes ATP turnover, allowing us to model the turbo design.

**Figure 1:**
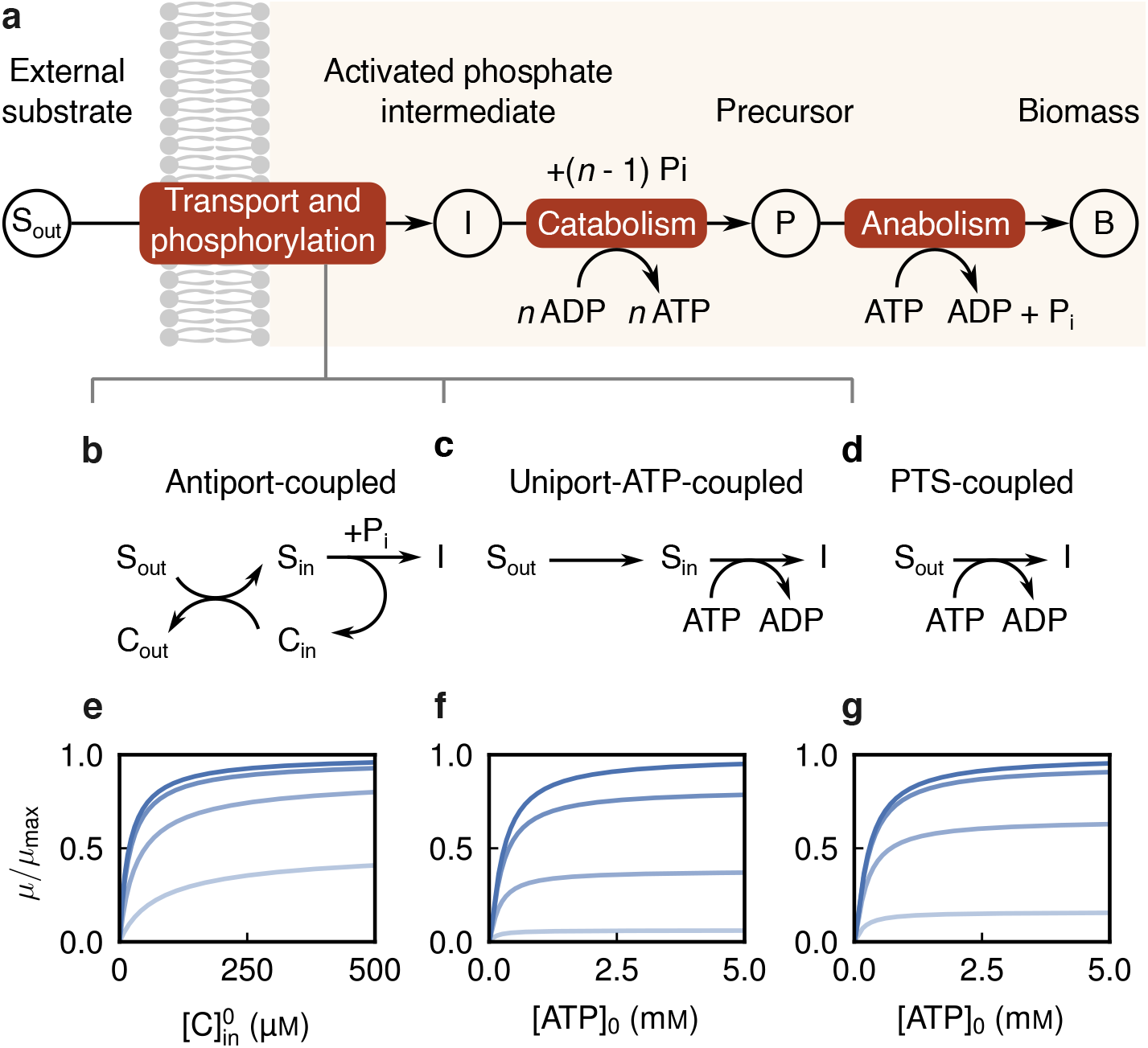
Cellular metabolism modeled as a series of lumped reactions. **a**, Schematic overview of the model. Here, “transport and phosphorylation” is used in a generic sense to denote the uptake and activation steps of the substrate, irrespective of whether these occur in separate reactions (**b, c**) or are mechanistically coupled in a single system such as the PTS (**d**). **b**, Antiport of the external substrate S_out_ for internal countersubstrate C_in_ coupled with phosphorylation of S_in_, which regenerates C_in_. For this mechanism, *n* = 1 (see catabolism). **c**, Uniport of the external substrate followed by ATP-driven phosphorylation of the internal substrate. For this mechanism, *n* = 2. **d**, Simultaneous transport and phosphorylation powered by ATP investment (PTS, phosphotransferase system). For this mechanism, *n* = 2. We set *n* = 2 to represent glycolysis-like activation, while noting that alternative pathways (e.g., Entner–Doudoroff) would correspond to different *n* values without altering the qualitative behavior. **e**–**g**, Normalized (specific) growth rate as a function of the initial C_in_ concentration in the antiport model (**e**); the initial ATP concentration in the uniport-ATP model (**f**); and the initial ATP concentration in the PTS model (**g**). Curves correspond to external substrate concentrations [S]_out_ of 1 µM, 10 µM, 0.1 mM, and 100 mM, shown as progressively darker shades of blue.

We examined three representative mechanisms for nutrient transport and phosphorylation, each implemented as a distinct cell model. The first mechanism, based on the ADI pathway and implemented as the antiport-coupled model (Fig. 1b), couples substrate antiport to phosphorylation, a reaction that both activates the incoming substrate [S]_in_ and regenerates the internal countersubstrate [C]_in_, thereby sustaining antiport. In this model, the sum [S]_in_ + [C]_in_ is constant and remains equal to its value at metabolic startup, 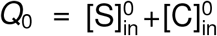. The second mechanism, typified by glycolysis in eukaryotes and implemented as the uniport-ATP-coupled model (Fig. 1c), involves facilitated diffusion of the external nutrient followed by ATP-driven phosphorylation in the cytosol. The third mechanism, used in the bacterial phosphotransferase system and implemented as the PTS-coupled model (Fig. 1d), combines transport and phosphorylation into a single phosphoenolpyruvate (PEP, equivalent of ATP)-driven reaction.^17^ In both ATP-coupled mechanisms, the sum [ADP] + [ATP] is constant and equal to *Q*_0_ = [ADP]_0_ + [ATP]_0_.

For each mechanism, the four above-mentioned reactions are modeled using Michaelis-Menten kinetics or the common modular rate law^18, 19^ for multisubstrate reactions. The rate of reaction *r* ∈ {t, p, c, a} is

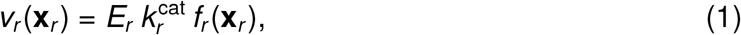

where *E*_*r*_ is the enzyme concentration, 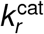 is the catalytic rate constant, and *f*_*r*_ is a function of the vector of metabolite concentrations **x**_*r*_. For instance, in the antiport-coupled model, 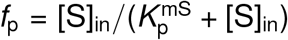, where 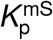 is the Michaelis constant. At steady state, each reaction carries the same flux *v*_*r*_ = *J*. Thus, using equation (1), we can express *E*_*r*_ in terms of *J* and **x**_*r*_, yielding the total enzyme concentration required for the cell to sustain the flux *J*:

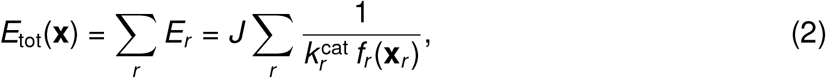

where **x** denotes the set of all metabolite concentrations involved in the four reactions.

To compute the steady-state growth rate of our cell model with the external substrate con-centration fixed, we proceeded as follows. First, we assume that a cell maximizes its specific growth rate and that it achieves this by optimizing its proteome *E*_*r*_ and internal metabolite concentrations **x**_in_, where **x**_in_ denotes the set of all internal metabolites in **x**. In contrast, the external substrate concentration [S]_out_ is determined by the surrounding environment and remains fixed. Second, we assume that the specific growth rate is proportional to the biomass production flux divided by the cellular biomass concentration. In our model, we approximate the biomass concentration by the total enzyme concentration, such that *µ* ∝ *J/E*_tot_, reflecting that proteins constitute the dominant fraction of cellular dry mass.^20^ Third, computationally, we fix the flux *J* and determine the minimal total enzyme concentration required to sustain this flux. Under these assumptions, maximizing the specific growth rate is equivalent to minimizing *E*_tot_ for a given flux, which corresponds to the Enzyme Cost Minimization principle.^21^

Unconstrained minimization of the total enzyme concentration, however, would drive some metabolite concentrations to unrealistically high values, because increasing such concentrations lowers *E*_tot_ (for details, see Methods). As intracellular metabolite pools cannot increase indefinitely without compromising cytoplasmic homeostasis (e.g., pH, ionic strength, and osmotic pressure), we impose a constraint on the total internal metabolite pool, 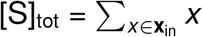. To enforce this constraint, we define the Lagrangian ℒ_*λ*_ with Lagrange multiplier *λ* as

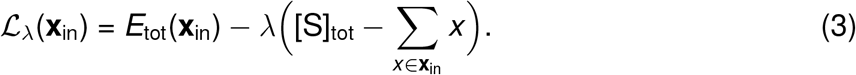

Minimizing the Lagrangian by solving *∂*ℒ_*λ*_*/∂***x**_in_ = 0 yields the optimized metabolite and enzyme concentrations. Substituting these concentrations into equation (2), and using *µ* ∝ *J/E*_tot_, we obtain the steady-state growth rate as a function of [S]_out_ and *Q*_0_. Full mathematical derivations are provided in the Methods.

### Growth depends on the initial Gibbs energy

To test whether growth is constrained by the initial available Gibbs energy, we asked whether cells starting with different internal metabolite concentrations—but placed in identical external conditions—achieve different steady-state growth rates. We tested this by computing steady-state growth rates for four values of the external substrate concentration [S]_out_ while systematically varying the initial internal concentrations. In the antiport-coupled model, we varied the internal countersubstrate concentration 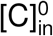 which modifies the chemical gradient across the membrane and thus the available Gibbs energy. In the ATP-coupled models, we varied the initial ATP concentration [ATP]_0_, which determines the Gibbs energy that can be released through ATP hydrolysis.

Consistent with our hypothesis, we found that varying 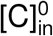or [ATP]_0_ produced substantial differences in steady-state growth rates at fixed [S]_out_ across all three models (Figs. 1e-1g). At high initial internal metabolite concentrations, cells reached their maximal specific growth rate, *µ*_max_, whereas at lower initial concentrations only limited growth rates were possible. This pattern was observed in all models, indicating that the limitation is independent of the underlying mechanism. Therefore, regardless of how substrate uptake is coupled to phosphorylation, cells starting with different internal metabolite concentrations in the same external environment achieve distinct steady-state growth rates, suggesting that growth is indeed constrained by the initial available Gibbs energy.

### Quantifying accessible Gibbs energy of metabolic activation

To establish a thermodynamic description of metabolic activation in the cell model, we derived expressions for the accessible Gibbs energy stored in metabolite configurations. In general, the combined cell-plus-environment system stores Gibbs energy, for example in transmembrane electrochemical gradients or intracellular ATP. However, not all of this energy is accessible, because metabolite concentrations are subject to physicochemical constraints and cannot be reorganized arbitrarily. We define the accessible Gibbs energy, denoted E, as the part of the total Gibbs energy that can be harnessed under such constraints. During metabolic activation, part of the initial accessible Gibbs energy E_0_ is dissipated as reaction fluxes increase toward their steady-state values. Once steady state is reached, the system retains a residual accessible Gibbs energy, which we denote ℰ_ss_. The difference Δℰ = ℰ_0_−ℰ_ss_ therefore quantifies the Gibbs energy dissipated to establish the steady state. For each cell model, we obtained expressions for E by integrating the Gibbs energy over the feasible concentration ranges permitted by the model’s physicochemical constraints (see Methods).

Towards examining how accessible Gibbs energy limits steady-state behavior, we quantified this energy in the antiport-coupled model at low initial countersubstrate concentration 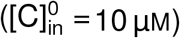. Specifically, we computed the initial and steady-state accessible Gibbs energies (equation (22), Methods) for different external substrate concentration [S]_out_ from very low (0.1 µM) to saturating (100 mM). At this low 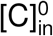, increasing [S]_out_ leads to only a modest increase in the initial accessible Gibbs energy E_0_ (Fig. 2a). Similarly, the dissipated energy Δ ℰ increases slightly with increasing [S]_out_, remaining below 5 · 10^*−*4^ kJ L^*−*1^ (shaded region in Fig. 2a). Thus, although the Gibbs energy change per mole of transported substrate, Δ*G*_antiport_ (equation (18), Methods), increases with [S]_out_, the accessible Gibbs energy hardly increases. This is because the limited initial countersubstrate concentration 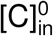 constrains the number of antiport exchanges that can occur, thereby restricting the Gibbs energy that can be dissipated. As a result, the accessible Gibbs energy remains bounded when 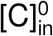 is low, even at high [S]_out_.

**Figure 2:**
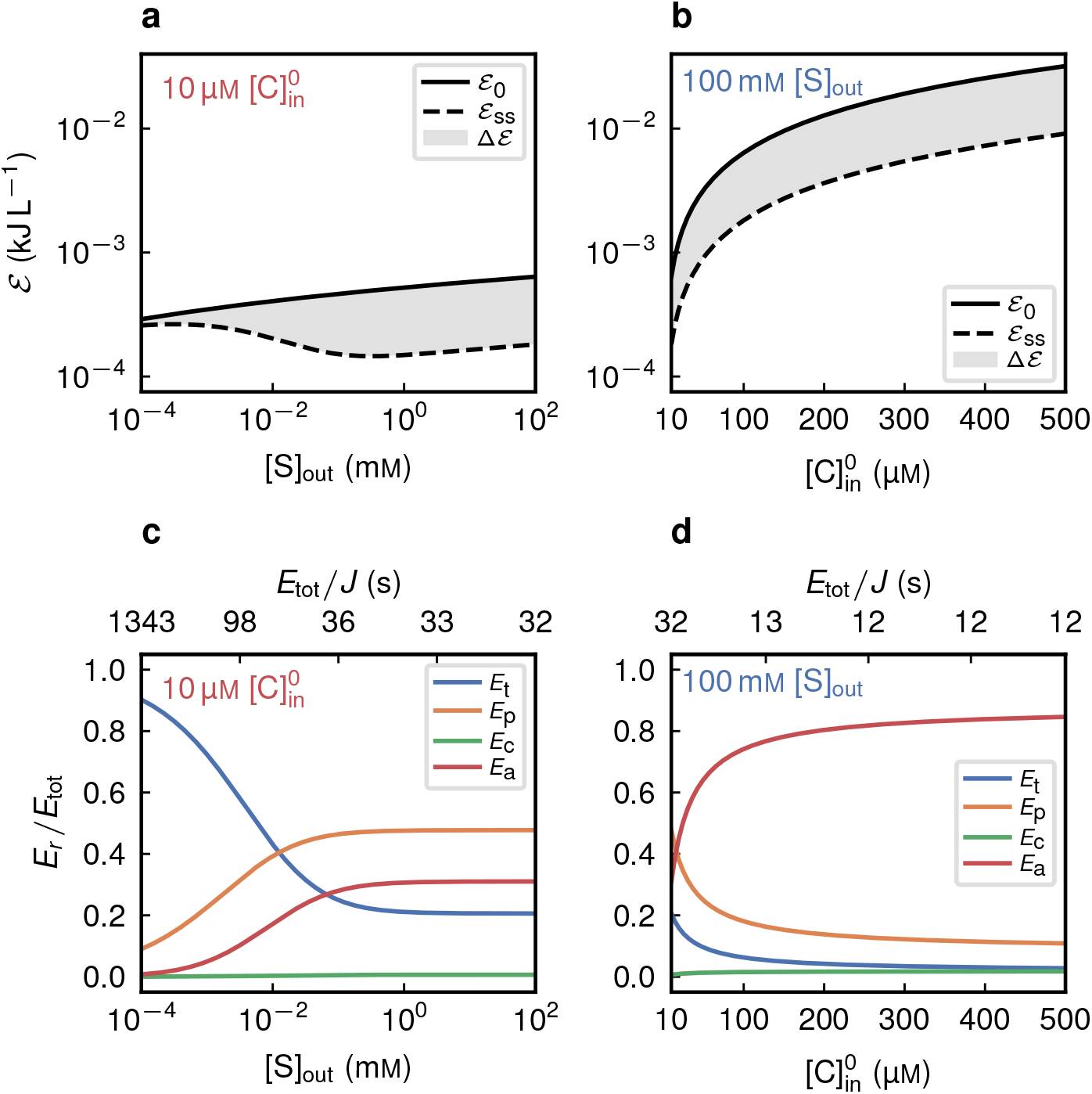
Accessible Gibbs energy limits metabolic activation in the antiport model. **a**, Initial accessible Gibbs energy (ℰ_0_, calculated from equation (22)), steady-state accessible Gibbs energy (ℰ_ss_), and dissipated energy Δ ℰ = ℰ_0_ − ℰ_ss_ as a function of the external substrate concentration [S]_out_, at a limiting initial internal countersubstrate concentration 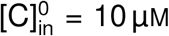 **c**, Proteome allocation to transport (*E*_t_), phosphorylation (*E*_p_), catabolic (*E*_c_), and anabolic (*E*_a_) enzymes, expressed as fractions of the total enzyme concentration *E*_tot_, for the simulations in panel **a**. The top axis is annotated with the corresponding total enzyme concentration per unit flux. **b** and **d**, Same analyses as in panels **a** and **c**, respectively, but as a function of 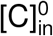 at saturating external substrate concentration [S]_out_ = 100 mM.

Next, we asked whether increasing the initial countersubstrate concentration increases the initial accessible Gibbs energy ℰ_0_. To this end, we simulated the antiport model at satu-rating [S]_out_ while varying 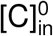. Indeed, as 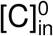 increased, ℰ_0_ rose substantially (Fig. 2b), and the dissipated energy Δ ℰ increased in parallel, reaching 2 · 10^*−*2^ kJ L^*−*1^ (shaded region in Fig. 2b). Together, these results show that the accessible Gibbs energy available for metabolic activation is strongly limited at low initial countersubstrate concentrations but can be substantially increased by elevating the initial countersubstrate concentration 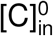.

### Accessible Gibbs energy constrains metabolic activation and steady-state growth rate

Since metabolite reorganization during metabolic activation is driven by the dissipation of accessible Gibbs energy, we asked how cells cope with limited accessible Gibbs energy in organizing their metabolite pools. We further asked whether sustaining a steady-state flux *J* under restricted metabolite reorganization requires compensatory proteome allocation, and whether this requirement could explain the observed growth constraints. To address these questions, we analyzed enzyme allocations normalized to total enzyme concentration, *E*_*r*_ */E*_tot_, for the two simulations discussed above.

In the first simulation (initial countersubstrate concentration 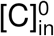 low), at very low external substrate concentration [S]_out_, the proteomic fraction of transporters is very high (Fig. 2c), consistent with several proteomic studies,^22–24^ and the total enzyme concentration per unit flux is high as well (upper horizontal axis of Fig. 2c). This proteome allocation reflects the need for large transporter concentrations to compensate weak transmembrane gradients at low [S]_out_ and 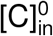. As the external substrate increases, both the transporter fraction and *E*_tot_ per flux decline, while the fraction of phosphorylation enzymes rises. The latter occurs because, at low 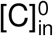, the attainable steady-state concentration [S]_in_ is low, so sustaining the flux *J* requires a larger proteome allocation to phosphorylation enzymes. As a result, the proteomic allocation to the investment enzymes (i.e., transport plus phosphorylation) is high across all external substrate levels, with the environment determining how this allocation is divided. In the second simulation (high initial 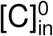), proteome allocation shifted: the fraction of investment enzymes (*E*_t_ + *E*_p_)*/E*_tot_ and the total enzyme concentration required per unit flux decreased, while the fraction of payoff enzymes (*E*_c_ + *E*_a_)*/E*_tot_ increased (Fig. 2d).

These results illustrate how cells cope with limited accessible Gibbs energy in the antiport-coupled model. When the initial accessible Gibbs energy ℰ_0_ is low, the metabolite concentrations can only be reorganized within narrow bounds set by the model’s physicochemical constraints. Consequently, sustaining a given steady-state flux *J* requires a compensatory increase in total enzyme concentration *E*_tot_ and larger proteomic fractions allocated to the investment enzymes (*E*_t_ and *E*_p_). Because growth rate scales as *µ* ∝ *J/E*_tot_, increasing *E*_tot_ lowers *µ*. In contrast, increasing the initial countersubstrate concentration raises the initial accessible Gibbs energy ℰ_0_, allowing greater dissipation of Gibbs energy toward metabolite reorganization. This redistribution of intracellular metabolite pools reduces the enzyme concentrations required to sustain flux, particularly for transport and phosphorylation reactions. The resulting decrease in *E*_tot_ allows the specific growth rate to approach its unconstrained maximum, *µ*_max_. In this regime of high accessible Gibbs energy, growth is controlled primarily by external substrate availability, consistent with Monod-like behavior.

To assess whether the growth limitation arising from a limited accessible Gibbs energy is a general feature of turbo-designed metabolism, we performed analogous simulations for the uniport-ATP- and PTS-coupled models. These simulations yielded behavior qualitatively similar to that of the antiport model. In the uniport-ATP model, low [ATP]_0_ limits the fraction of substrate that can be phosphorylated, reducing the initial accessible Gibbs energy ℰ_0_ (Supplementary Fig. 1a) and increasing the fraction of phosphorylation enzymes required to sustain steady-state flux (Supplementary Fig. 1c). In the PTS model, low [ATP]_0_ directly constrains substrate uptake, resulting in a higher fraction of transport enzymes (Supplementary Fig. 2c). Consistently, increasing [ATP]_0_ in both models increases ℰ_0_ and the dissipated energy Δ ℰ (Supplementary Figs. 1b and 2b), enabling reallocation of the proteome, a reduction in the total enzyme concentration *E*_tot_ per unit flux, and growth rates approaching the unconstrained maximum, *µ*_max_ (Supplementary Figs. 1d and 2d). Together, these results confirm that growth limitation by accessible Gibbs energy, and the associated proteomic burden of the investment reactions, is a general feature of the turbo-designed metabolic pathways considered.

Although our models indicate that cells cope with limited accessible Gibbs energy through coordinated organization of both metabolite pools and proteome allocation, we sought to isolate the contribution of metabolite reorganization (in contrast to proteome allocation) by analyzing a scenario in which enzyme levels were held fixed. Using the antiport-coupled model, we first determined the enzyme allocation under saturating external substrate ([S]_out_ = 100 mM) and high initial countersubstrate concentration 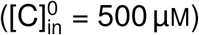. We then fixed these enzyme levels and progressively decreased 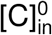, computing at each value the maximal steady-state flux *J* compatible with the fixed proteome. Because growth scales as *µ* ∝ *J/E*_tot_, and *E*_tot_ was held constant in this analysis, any reduction in growth must arise from a decrease in *J*. Indeed, growth rates declined with decreasing 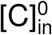 (Supplementary Fig. 3), demonstrating that a reduction in accessible Gibbs energy is sufficient to limit steady-state growth even in the absence of the possibility for proteome reallocation. Quantitatively, growth under fixed proteome remained below that achieved with full optimization, indicating that proteome reallocation expands the achievable flux but does not remove the underlying energetic constraint. These results show that metabolite reorganization and proteome reallocation are complementary responses to a limited accessible Gibbs energy.

### Experimental evidence for energy-dependent constraint on growth

While our coarse-grained model of cellular metabolism suggests that the specific growth rate is limited by accessible Gibbs energy, an important question is whether this dependence extends to real biological cells. However, because the limitations on metabolic reorganization predicted by our model emerge from system-level coupling between transport, metabolism, and biosynthesis, experimental validation *in vivo* is challenging. We therefore sought experimental evidence from a minimal, controllable system that captures the features of turbo-designed metabolism. To this end, we reconstituted the arginine deiminase (ADI) pathway in synthetic vesicles.^25–30^ These vesicles allow experimental control over enzyme concentrations and initial metabolite pools, enabling us to test whether differences in the initial accessible Gibbs energy translate into differences in long-term metabolic capacity.

The ADI pathway is composed of the arginine-ornithine antiporter (ArcD) and three cytosolic enzymes (Fig. 3a). In this pathway, external arginine is exchanged by intravesicular ornithine (ArcD). Then, intravesicular arginine is hydrolyzed by arginine deiminase (ArcA), forming citrulline and NH_4_^+^. Citrulline is then converted, along with P_i_, into carbamoyl phosphate (CP) and ornithine by ornithine transcarbamoylase (ArcB). Finally, carbamate kinase (ArcC) catalyzes the reaction between CP and MgADP to produce MgATP and carbamate. To mimic catabolism with downstream ATP consumption, analogous to ATP usage for biomass production in cells, we added the mitochondrial ATP-ADP carrier (AAC)^27^ to the vesicles. This protein exports synthesized ATP in exchange for external ADP, which is equivalent to ATP consumption by hydrolysis.

**Figure 3:**
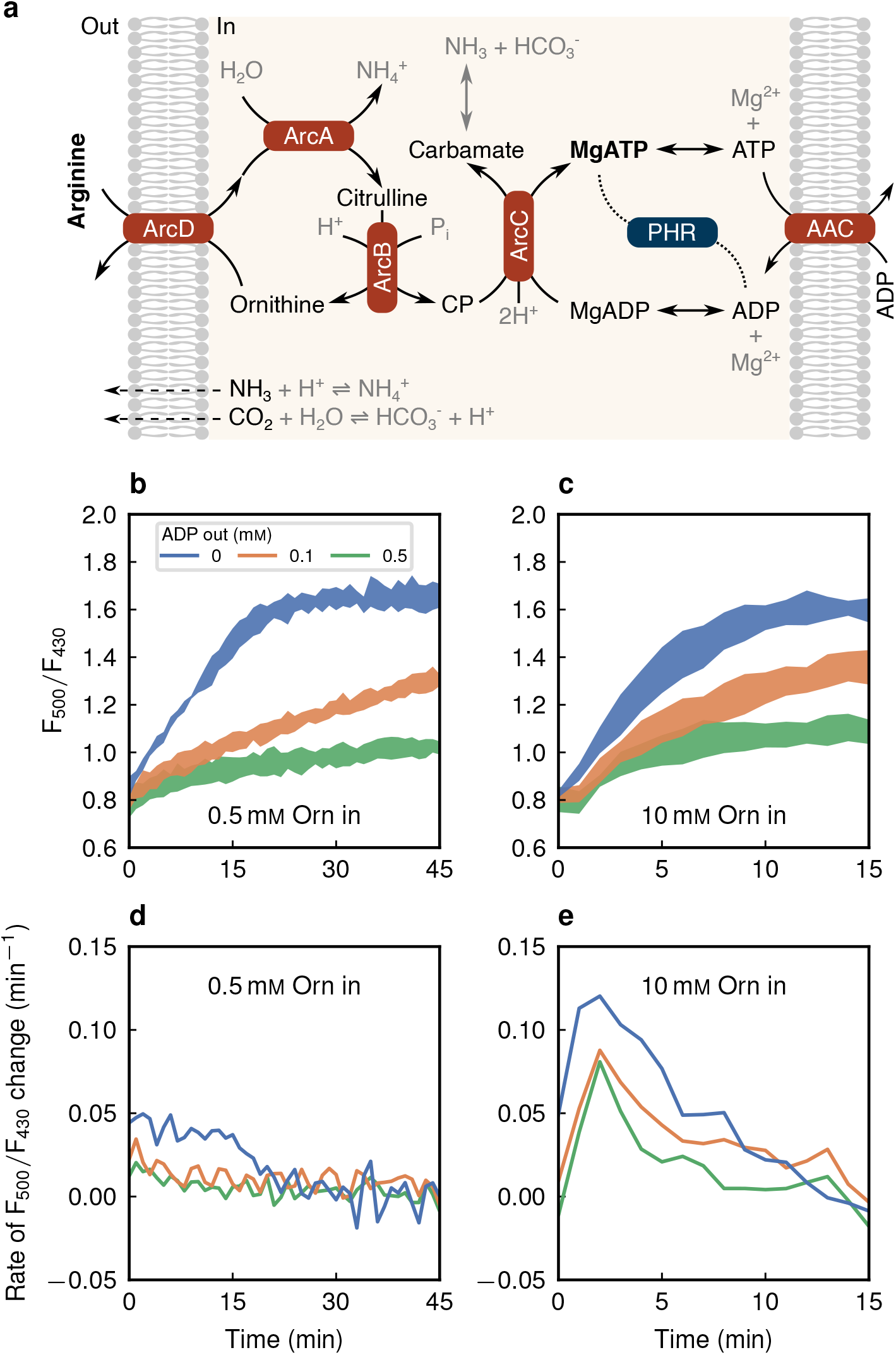
Higher ornithine inside drives faster ATP production despite identical arginine outside. **a**, Schematic representation of the arginine breakdown pathway. **b** and **c**, PercevalHR ratiometric signal (F_500_*/*F_430_) for the ADI pathway reconstituted in synthetic vesicles. Data represent the mean of biological triplicates, with the shaded area indicating ± one standard deviation. The system was initialized with either 0.5 mM (**b**) or 10 mM (**c**) ornithine inside and three concentrations of ADP outside. **d** and **e**, Instantaneous rate of change of F_500_*/*F_430_ calculated from the experimental traces in **b** and **c**.

We prepared vesicles by encapsulating the cytosolic enzymes of the ADI pathway, with the arginine-ornithine antiporter ArcD and the ATP-ADP carrier AAC reconstituted in the membrane. These vesicles were exposed to a saturating concentration of external arginine (10 mM) and to three different external ADP levels: none (to prevent ATP export via the carrier, hence no ATP consumption), 0.1 mM, or 0.5 mM. For each condition, the only parameter varied was the initial intravesicular ornithine concentration, set to either low (0.5 mM) or high (10 mM), creating two markedly different initial accessible Gibbs energies while keeping all other system parameters identical. To dynamically monitor the vesicles’ energetic state, we co-encapsulated the ratiometric fluorescent probe PercevalHR and tracked the fluorescence ratio F_500_*/*F_430_, a proxy for the intravesicular ATP-to-ADP ratio^31^ (Supplementary Fig. 9).

In these experiments, we first examined the time courses of the PercevalHR ratio after external arginine addition in the absence of external ADP. Over the course of a few minutes, the ratios rose and saturated (Figs. 3b and 3c). While both initial ornithine conditions reached similar final PercevalHR ratios, the speed at which this ratio was achieved differed markedly: in vesicles with 10 mM ornithine, the ratio increased 2.4× faster than in those with 0.5 mM ornithine. Consistently, the rate of change of F_500_*/*F_430_, obtained by numerically differentiating the experimental trace, was markedly higher at 10 mM ornithine (Figs. 3d and 3e). These differences persisted when external ADP was added. Here, although the final PercevalHR ratio was lower in the presence of external ADP (due to ATP export via AAC), it was reached over a similar timescale and with comparable rates. These results demonstrate that the initial intravesicular ornithine concentration determines how rapidly the ADI pathway can elevate the ATP-to-ADP ratio.

To connect these observations to steady-state growth rate, we first noted that, under conservation of the total nucleotide concentration, the rate of change of the PercevalHR fluorescence ratio F_500_*/*F_430_ is proportional to the net ATP production rate of the pathway (equation (32), Methods). Therefore, the markedly higher rate of change observed with 10 mM initial ornithine reflects a higher ATP production rate and flux through the entire pathway. Thus, vesicles initiated with higher ornithine, and therefore higher initial accessible Gibbs energy, are expected to achieve and sustain a higher steady-state growth rate once coupled to growth processes. These vesicle experiments provide experimental evidence that the energetic state at metabolic activation imposes a constraint on long-term ATP production rate and, by extension, on long-term growth rate.

### Citrulline accumulation mediates control of steady-state ATP production rate

To investigate how different steady-state ATP production rates arise from initial metabolite levels and their dynamics during activation, we constructed and analyzed a kinetic model of the ADI pathway. To this end, we measured transport and enzymatic activities in isolation, and fitted comprehensive rate laws to the data. ArcA (Fig. 4a) and ArcB (Figs. 4b and 4c) activities were measured by HPLC and spectroscopic readout, ArcC activity via ATP luminescence assays (Fig. 4d), and ArcD-mediated transport using radiolabeled arginine with ornithine as countersubstrate (Figs. 4e and 4f). The resulting rate laws were integrated into an ODE-model describing the entire pathway. The few remaining parameter values that could not be measured in isolation were obtained by fitting the model to the experimentally measured F_500_*/*F_430_ ratios shown in the previous section. Details of the model construction, parameter estimation, and experimental measurements are provided in the Methods. The fitted model reproduces these measurements with good agreement (Supplementary Fig. 10).

**Figure 4:**
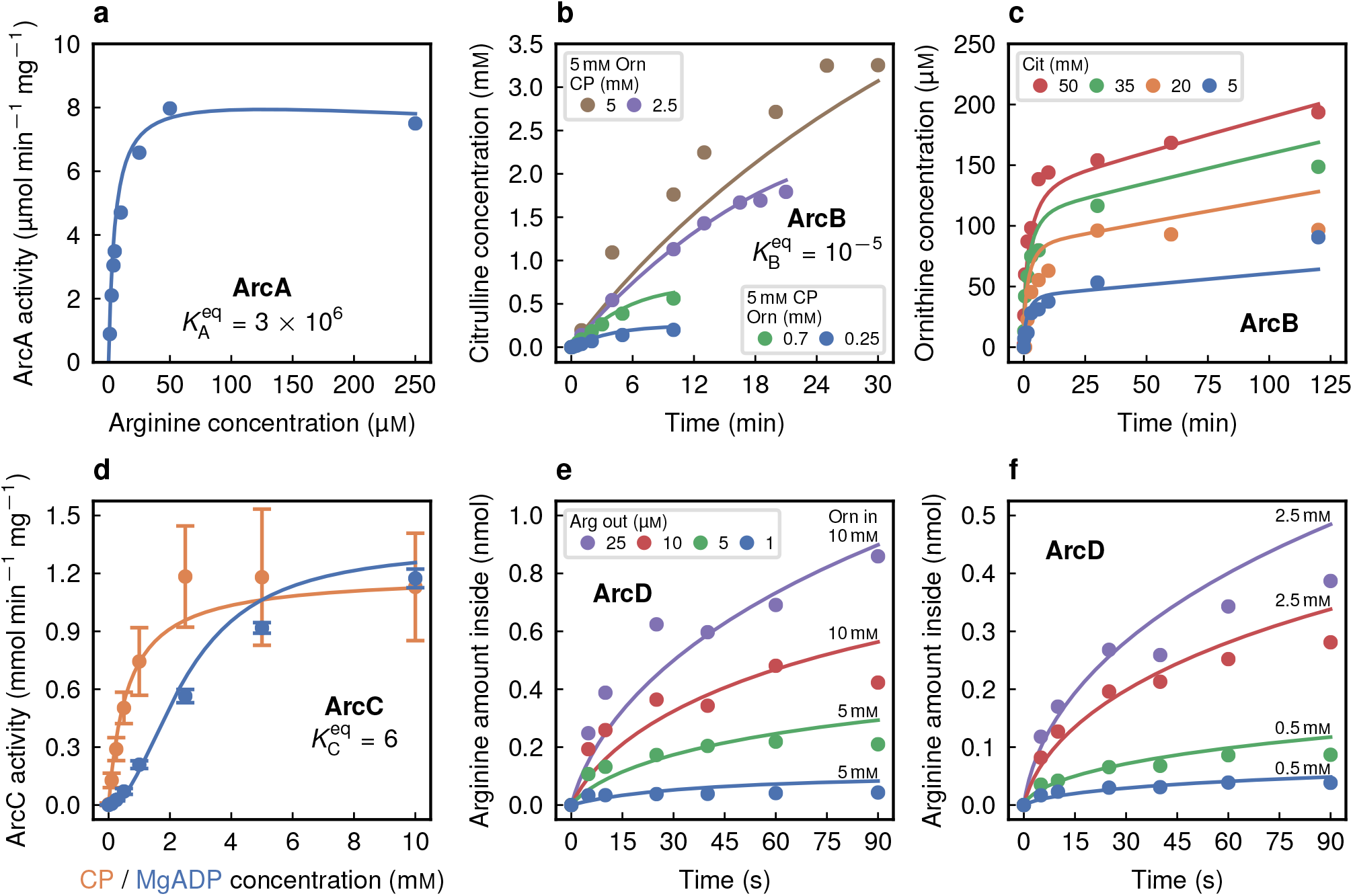
Characterization of the antiporter and the three cytosolic enzymes that constitute the arginine breakdown pathway. **a**, Arginine deiminase (ArcA) characterization. Michaelis-Menten plot of ArcA activity, defined as the initial rate of citrulline formation per milligram of ArcA, versus initial arginine concentration. **b**, Ornithine transcarbamoylase (ArcB) characterization in the thermodynamically favorable direction. Dynamic citrulline formation under selected initial conditions. For the complete dataset, please refer to Supplementary Fig. 5. **c**, Ornithine transcarbamoylase (ArcB) characterization in the thermodynamically unfavorable direction. Dynamic ornithine formation. **d**, Carbamate kinase (ArcC1) characterization in the thermodynamically favorable direction. **e** and **f**, Arginine-ornithine antiporter (ArcD) characterization. Dynamic arginine uptake in proteoliposomes under selected initial conditions. For the complete dataset, please refer to Supplementary Fig. 8.

Using this kinetic model, we investigated the transient behavior of internal metabolite pools in the ADI pathway until steady state is reached. Our experiments (cf. Fig. 3) showed that the initial intravesicular ornithine concentration influences the ATP production rate. We then tested whether ornithine directly controls the steady state or acts indirectly through another variable. As a candidate, we examined the total conserved metabolite pool 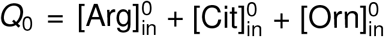 Simulations at *Q*_0_ = 0.5 mM starting from four distinct initial conditions— only ornithine 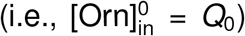, only citrulline, only arginine, or an equimolar mixture of the three—converged to the same steady-state metabolite composition (Fig. 5a), despite markedly different transient dynamics (Supplementary Figs. 11a–11c). This metabolite composition substantially shifted when we increased *Q*_0_ to 10 mM (Supplementary Figs. 11d– 11f). Together, these results demonstrate that the steady-state metabolite composition depends on the total pool *Q*_0_, but is independent of the initial distribution among arginine, citrulline, and ornithine.

**Figure 5:**
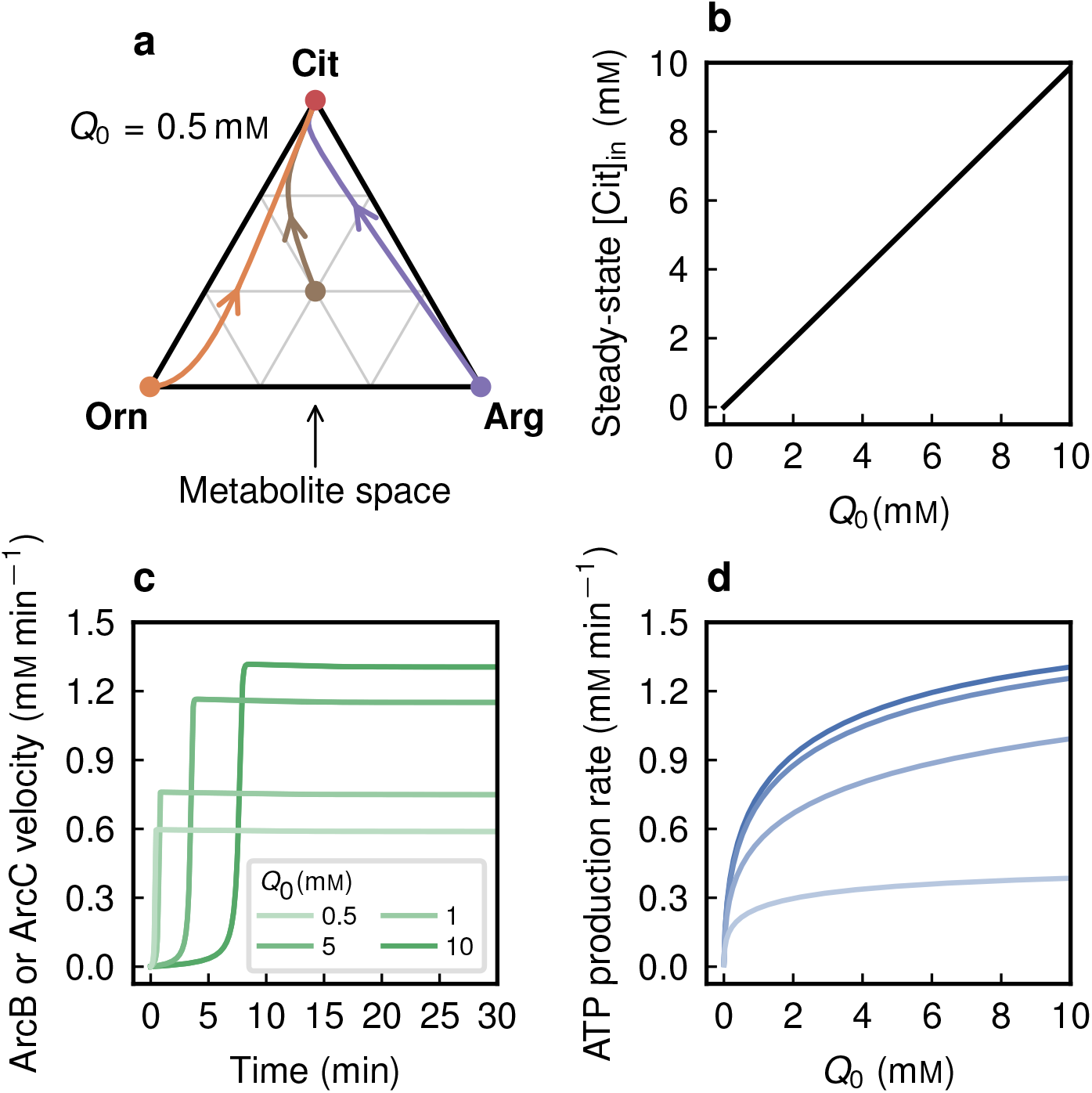
Dynamic intravesicular metabolite reorganization links the initial conditions to the resulting steady-state ATP production rate. **a**, Ternary plot showing the time evolution in metabolite space of intravesicular arginine, citrulline, and ornithine concentrations at fixed *Q*_0_ = 0.5 mM and fixed external arginine and ADP concentrations (100 mM), for four initial conditions: only arginine 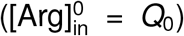, only citrulline 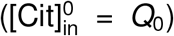, only ornithine 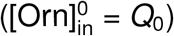 and an equimolar mixture of the three 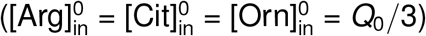 Filled circles indicate the initial conditions, and arrows indicate the direction of time evolution. **b**, Steady-state intravesicular citrulline concentration as a function of *Q*_0_. **c**, Time evolution of ArcB and ArcC reaction velocities (overlapping) at fixed external arginine and ADP concentrations (100 mM), for the same four values of *Q*_0_ shown in **b. d**, Steady-state ATP production rate, obtained from the long-time limit of the simulations, as a function of *Q*_0_. Curves correspond to external arginine concentrations of 1 µM, 10 µM, 0.1 mM, and 100 mM, shown in progressively darker shades of blue. Across all conditions, the external ADP concentration was held at 100 mM.

Next, we analyzed how the steady-state metabolite concentrations determined by *Q*_0_ affects the ATP production rate. While intravesicular arginine and ornithine relax to low steady-state concentrations, citrulline accumulates to levels comparable to *Q*_0_ (Fig. 5b), so that steady-state citrulline concentration increases with *Q*_0_ and encodes the size of the initial metabolite pool. This accumulation has direct kinetic consequences: since the ArcB reaction rate increases with citrulline concentration (equation (24)), higher citrulline levels lead to increased ArcB flux (Fig. 5c). Because the pathway operates under quasi-steady-state conditions before reaching a final steady-state (Fig. 5c), the flux through ArcB sets the flux through ArcC, and thus the ATP production rate. In this way, citrulline acts as the intermediate variable that transmits the effect of *Q*_0_ to steady-state ATP production, with higher *Q*_0_ enabling higher steady-state ATP production rates.

In agreement with the coarse-grained antiport-coupled model (Fig. 1e), the steady-state ATP production rate saturates with increasing *Q*_0_ (Fig. 5d). For a fixed *Q*_0_, ATP production rises with external arginine, showing control by the internal pool *Q*_0_ and external substrate. This dual dependence parallels that of accessible Gibbs energy on *Q*_0_ and external arginine (equation (22), Methods). In the kinetic model, *Q*_0_ sets the steady-state citrulline concentration, while external arginine drives ornithine export via ArcD, sustaining net forward flux through the thermodynamically unfavorable ArcB reaction. At large *Q*_0_, the ATP production rate is controlled primarily by external arginine (flat region in Fig. 5d), recovering the classical Monod regime.

## Discussion

Activating metabolism after starvation is commonly viewed as a process that restores steady-state growth once nutrients become available. Here, we show that this view is incomplete. Using coarse-grained models of turbo-designed metabolic pathways, we demonstrate that metabolic activation requires an upfront energetic investment drawn from the accessible Gibbs energy at activation, which in turn constrains the achievable steady-state growth rates. This history dependence arises because, in turbo-designed pathways, the accessible Gibbs energy is not set by external nutrient availability alone but also by the initial internal metabolite pools. The accessible Gibbs energy available at activation then limits the extent of possible metabolic reorganization, thereby constraining the long-term metabolic capacity. We validated this prediction using vesicles encapsulating the arginine deiminase pathway, where long-term ATP production rate depends on the conserved pool of arginine, citrulline, and ornithine via citrulline accumulation, which links the initial accessible Gibbs energy to long-term metabolic output. Together, these results establish metabolic activation as a process that determines steady-state cellular growth, extending beyond the memory-less assumptions of classical growth laws such as the Monod equation and its extensions (e.g., the Contois and Teissier models^33, 34^). These models emerge as special cases when the initial accessible Gibbs energy is abundant.

Our results suggest that the constraints linking initial accessible Gibbs energy to long-term cell growth emerge through a series of interdependent steps (Fig. 6). Nutrient exposure and the initial metabolite pool configuration set the initial accessible Gibbs energy (ℰ_0_). During metabolic activation, part of this energy is irreversibly dissipated (Δℰ), driving reorganization of the internal metabolite pool. However, the finite dissipated Gibbs energy Δℰ limits metabolite reorganization. In the ADI pathway vesicle system, where enzyme concentrations are fixed, this limited metabolite reorganization sets the maximal steady-state flux *J* under the given proteome *E*_tot_; growth rate then follows from *µ* ∝ *J/E*_tot_. In contrast, in the cell model (mimicking living cells where enzyme concentrations vary), constrained reorganization sets the *E*_tot_ required to sustain a given *J*, with *µ* ultimately bounded by steady-state metabolite and proteome organization. Thus, by restricting the metabolic states that can be reached at steady state, the initial accessible Gibbs energy generates a form of metabolic memory that manifests as a constrained steady-state growth rate.

**Figure 6:**
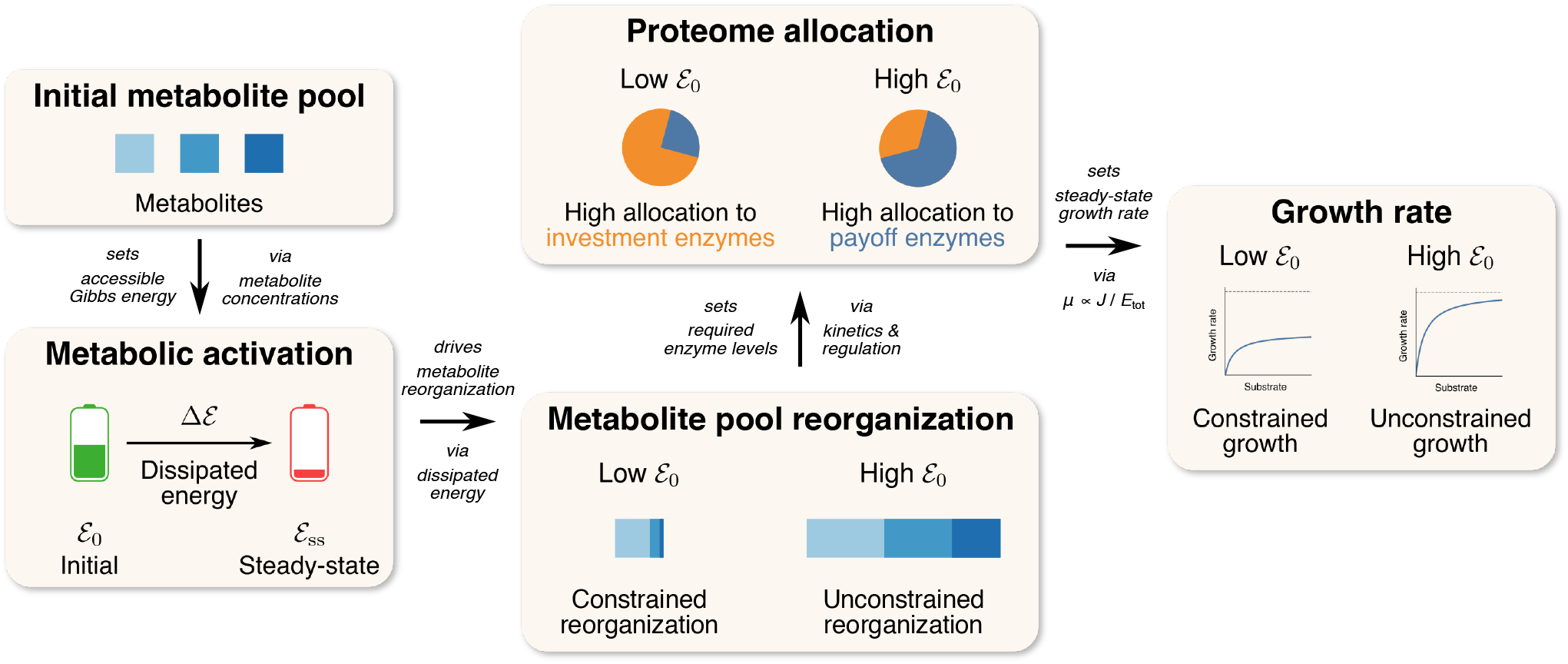
How a cell’s initial accessible Gibbs energy constrains its long-term growth rate. Upon nutrient exposure, the cell begins from an initial metabolite pool configuration (e.g., external arginine and internal ornithine in reconstituted ADI vesicles). This configuration *sets* the initial accessible Gibbs energy (ℰ_0_). During metabolic activation, part of this energy is irreversibly dissipated (Δℰ). The dissipated energy *drives* reorganization of the internal metabolite pool; however, because Δℰ is finite, it limits the extent to which intracellular metabolites can be reorganized (e.g., the steady-state citrulline concentration is low when ℰ_0_ is low). When enzyme concentrations are fixed (as in vesicles), metabolite reorganization alone *sets* the maximal steady-state flux *J* that the pathway can sustain under the given fixed proteome *E*_tot_. Specific growth rate then follows from *µ* ∝*J/E*_tot_. In contrast, when enzyme concentrations can vary (as in living cells), metabolite reorganization *sets* the total enzyme concentration *E*_tot_ required to sustain a given flux *J*. The steady-state flux *J* together with the total enzyme concentration *E*_tot_ (here, an approximation for biomass), both set by metabolite and proteome steady-state organization, *sets* the specific growth rate *µ* ∝ *J/E*_tot_. In this way, the initial accessible Gibbs energy constrains the region of metabolic states that can be reached and ultimately bounds the steady-state growth rate the cell can attain.

Our definition of accessible Gibbs energy is conceptually related to the thermodynamic notion of exergy,^35, 36^as both quantify the fraction of stored energy that can be converted into useful work under given constraints. Beyond this conceptual link, our definition maps directly onto the nonequilibrium Gibbs energy introduced by Rao and Esposito.^37^In their theory, the nonequilibrium Gibbs energy is determined by the conserved quantities of the reaction network; in our models, these correspond to conserved metabolite pools, *Q*_0_, which bound the extent of metabolite reorganization and thereby set the Gibbs energy available for dissipation during metabolic activation.

The amount of Gibbs energy that can be accessed at steady state (ℰ_ss_) is constrained by the energy dissipated (Δℰ) during metabolic activation. Therefore, accessible Gibbs energy is not determined solely by the instantaneous metabolite concentrations, but reflects the system’s prior trajectory. The metabolic memory we identify thus represents a biological instance of thermodynamic memory, as described in irreversible thermodynamics, where the state of a system retains signatures of past dissipation during transient processes.^38^Unlike classical memory encoded in gene regulation or signaling networks, this form of memory arises directly from the design of the pathway. In turbo-designed metabolic pathways, energy dissipated during activation irreversibly reduces the accessible Gibbs energy, making steady-state properties dependent on the system’s transient history. This history-dependent constraint parallels theoretical results by England, who showed that the minimal heat required for bacterial self-replication depends not only on the entropy change associated with doubling but also on the time over which replication occurs.^39^In both cases, Gibbs energy dissipation imposes limits on the system’s performance—growth rate in our work, replication efficiency in England’s—that depend on the transient history of the process.

Our finding that the accessible Gibbs energy at activation constrains long-term growth has important implications. For instance, to avoid the severe long-term consequences of a low steady-state growth rate after startup, it would be advantageous for cells experiencing starvation to secure a minimal level of accessible Gibbs energy. Indeed, persister cells formed after sudden nutrient shifts have been reported to maintain ATP levels despite strongly reduced metabolic activity.^40–42^ Furthermore, mechanisms such as toxin-antitoxin systems, which suppress energy-consuming processes under unfavorable conditions, likely contribute to preserving accessible Gibbs energy. By limiting dissipation during stress, such mechanisms may safeguard the capacity for metabolic activation and high growth rates when conditions improve.

More broadly, our work frames metabolic regulation as a problem of managing energy over time, in addition to optimizing steady-state fluxes. Our work provides a framework that may help interpret lag phases, persistence, and variability in microbial recovery, and offers potential guidance for designing synthetic metabolic systems that minimize costly activation bottlenecks.

## Methods

### Development of the antiport-coupled cell model

We constructed a cell model based on the framework introduced by Noor et al. ^16^The reactions were modeled using the common modular rate law.^18, 19^In the antiport-coupled cell model (Fig. 1b), we assumed that the external countersubstrate C_out_ was maintained at a constant, very low concentration (1 pM). Accordingly, antiport was modeled as

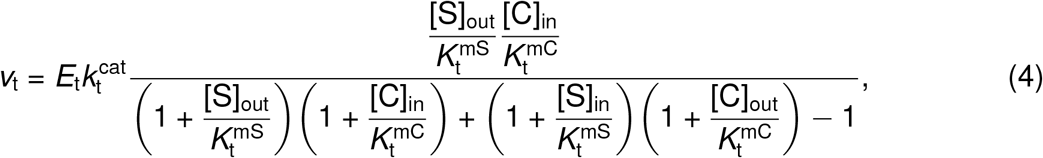

where *E*_t_ is the concentration of transporters, 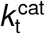is the catalytic rate constant, 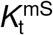 and 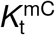 are Michaelis constants, and both external and internal (counter-)substrates were assumed to have equal affinity for their respective binding sites. Phosphorylation was modeled as incorporation of cytosolic P_i_ onto the internal substrate S_in_, with the concentrations of P_i_, ADP, and ATP assumed to remain fixed and non-limiting throughout metabolic activation. Phosphorylation of S_in_ was modeled as an irreversible reaction:

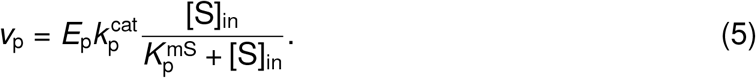

In catabolism, the P_i_ incorporated into the intermediate I is transferred to ADP, generating ATP and the precursor P. The catabolic reaction was modeled as

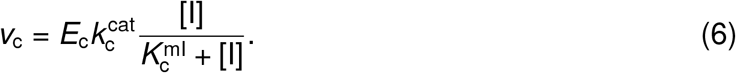

Anabolism was modeled as

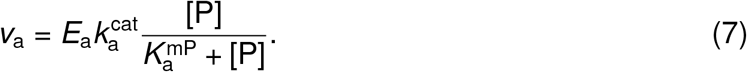

At steady state, each reaction carries the same flux *J*. Substituting this into Eqs. 4–7, we obtained expressions for the enzyme level of each reaction as a function of *J* and metabolite concentrations. Summing these levels gave the total enzyme concentration required to sustain the flux *J*:

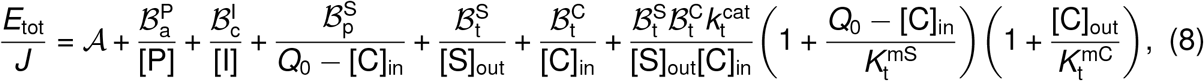

Where 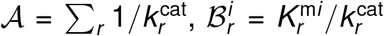and all metabolite concentrations are at steady state. The Enzyme Cost Minimization principle was adopted,^21^according to which growth rate was maximized by minimizing *E*_tot_ for a fixed flux *J*, with growth rate taken proportional to *J/E*_tot_. A constraint was imposed on the total pool of internal metabolites, as unconstrained minimization of *E*_tot_ would otherwise drive concentrations of I and P to unrealistically high values (equation (8)):

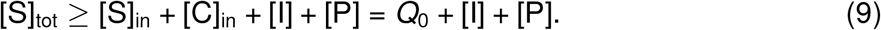

The constraint was enforced using Lagrange’s method. A Lagrangian function with Lagrange multiplier *λ* was defined as

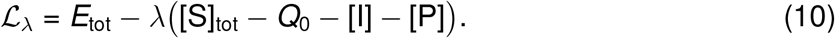

Minimizing ℒ_*λ*_ with respect to [I] and [P] yielded

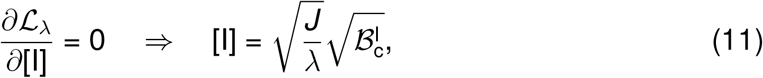

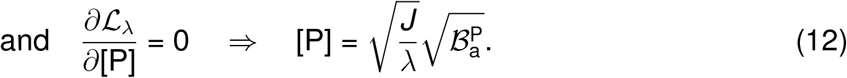

The Lagrange multiplier *λ* was determined by assuming the constraint was saturated, such that the total internal metabolite pool reached its upper bound (equation (9)). Solving for 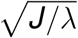and substituting back gave

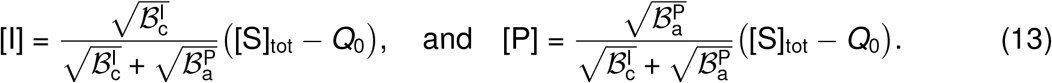

Minimizing the Lagrangian with respect to [C]_in_ yielded the quadratic equation

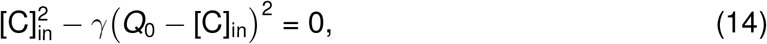

where *γ* is defined as

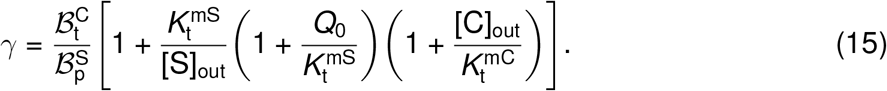

Although the quadratic has two roots, only one corresponds to a physically feasible concentration. The valid solution is

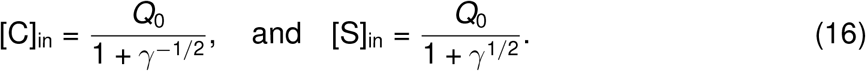

These expressions were substituted into equation (8) to compute the normalized growth rate *µ/µ*_max_, where *µ*_max_ ∝ 1*/*𝒜, as a function of the external metabolite concentrations and *Q*_0_.

For the simulations (Figs. 1e and 2), we used the catalytic rate constants 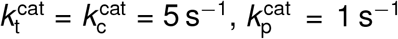, and 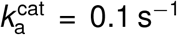, and the Michaelis constants 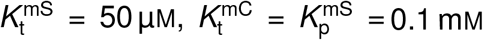, and 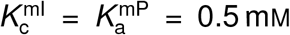. Additionally, we set [S]_tot_ = 100 mM, and considered 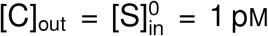,that is, essentially zero. The uniport-ATP- and PTS-coupled models were constructed following the same optimization framework, with full derivations and parameter values provided in Supplementary Methods.

### Quantifying accessible Gibbs energy of the antiport-coupled model

The antiport transport of an infinitesimal amount of substrate across the membrane requires an energy per unit volume

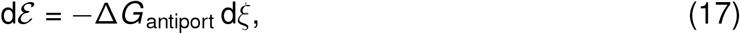

where d*ξ* denotes the change in intracellular substrate concentration associated with the transported amount, and Δ*G*_antiport_ is the Gibbs energy change per mole of exchanged substrate, given by

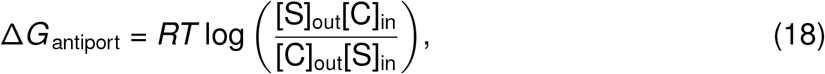

with *R* being the ideal gas constant and *T* the absolute temperature.

Consider a reference state defined by the intracellular concentrations 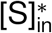 and 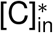, at which we want to compute the accessible Gibbs energy. Upon antiport transport, these concentrations change according to

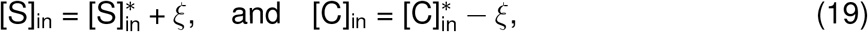

while the external concentrations [S]_out_ and [C]_out_ are held constant. Substituting these expressions into equation (17) yielded

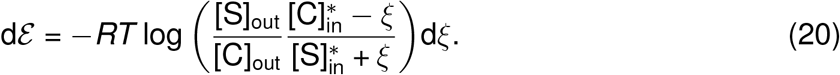

The accessible Gibbs energy was obtained by integrating dℰ up to the point at which no further Gibbs energy can be extracted from the substrate configuration, that is, up to the condition dℰ= 0. This condition defines the maximum achievable concentration change:

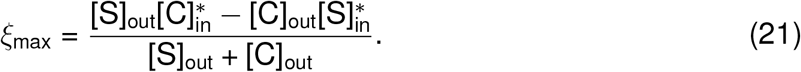

In the case [C]_out_ = 0, this reduces to 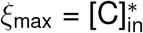, indicating that the maximum change in intracellular concentration is limited by the initial concentration of the internal countersubstrate.

Performing the integration and substituting the above expression for *ξ*_max_ gave a closed-form result for the accessible Gibbs energy:

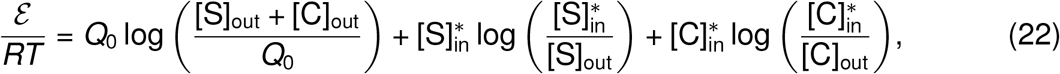

where we used that 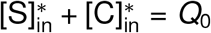. Expressions for the accessible Gibbs energies for the uniport-ATP- and PTS-coupled models were derived following the same procedure and are provided in Supplementary Methods.

### *L. lactis* arginine deiminase (ArcA) characterization

Arginine deiminase catalyzes the strongly favorable hydrolysis of arginine into citrulline and 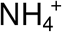. Enzymatic assays were performed with arginine as the substrate (Fig. 4a and Supplementary Fig. 4). Citrulline formation over time was quantified via RP-HPLC of the aminoenone derivative, with absorbance measured at 280 nm. L-Arginine was added at different concentrations to a reaction solution containing 42 nM purified ArcA and 50 mM KP_i_ buffer at pH 7.0, in a final volume of 700 µL. Reactions were run at 30 °C.

Substrate inhibition was previously reported for *Lactococcus lactis* ArcA, with a 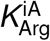 of 3.2 mM.^26^ ArcA activity was therefore modeled using the Michaelis-Menten equation with uncompetitive inhibition:

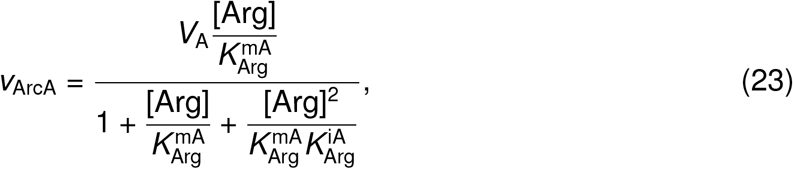

where 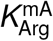 is the Michaelis constant, and *V*_A_ = 4[ArcA]*k*_A_ is the limiting velocity, [ArcA] being the enzyme concentration, and *k*_A_ the catalytic rate constant. The factor of 4 arises from the assumption that the tetramer ArcA has one active site per monomer.

The parameters of the rate law were obtained by minimizing the residual sum of squares between the HPLC data and the solution to the equation d[Cit]*/*d*t* = *v*_ArcA_ (Supplementary Table 1). The inhibition constant was fixed at its previously established value.

### *L. lactis* ornithine transcarbamoylase (ArcB) characterization

Ornithine transcarbamoylase catalyzes the conversion of citrulline and P_i_ into carbamoyl phosphate (CP) and ornithine. The apparent equilibrium constant of this reaction is 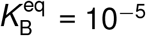 (Supplementary Table 1), indicating that ArcB operates in the thermodynamically unfavorable direction for ATP synthesis in the ADI pathway.

Assays were performed in both reaction directions (Figs. 4b and 4c and Supplementary Fig. 5). In the thermodynamically favorable direction, L-ornithine was added at different concentrations to a reaction solution containing 50 mM KP_i_ buffer at pH 7.0, 5 mM CP, and 15 nM ArcB. When CP dependence was tested, L-ornithine was kept constant at 5 mM. CP was added shortly before L-ornithine to minimize degradation during pre-incubation. Citrulline formation was quantified via RP-HPLC of the aminoenone derivative, with absorbance measured at 280 nm. In the thermodynamically unfavorable direction, citrulline was added at concentrations ranging from 5 to 100 mM to a reaction solution containing 50 mM KP_i_ buffer at pH 7.0 and 15 nM ArcB, and ornithine formation was quantified via RP-HPLC, with absorbance measured at 280 nm. All reactions were run at 30 °C. Given that assays were monitored for up to 120 min, CP instability was accounted for in the model. CP degradation was characterized by monitoring a 100 mM CP solution in 50 mM KP_i_ buffer at pH 7.0 by ^31^PNMR on a Varian Mercury Plus 300 MHz system using a Varian 5 mm PFG AutoSW probe. Data were collected for 812 min, with one spectrum recorded every 21 min over 256 scans (Supplementary Fig. 6), yielding a hydrolysis rate constant of 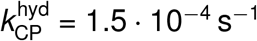.

ArcB follows an ordered bi-bi mechanism in which CP binds first, followed by ornithine, with citrulline released first and P_i_ second.^43–45^Therefore, ArcB activity was modeled using this mechanism with the steady-state approach:^32^

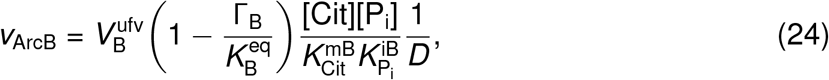

where 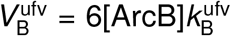 is the limiting velocity of the hexameric enzyme in the unfavorable direction, Γ_B_ is the mass-action ratio, and the denominator *D* is defined as

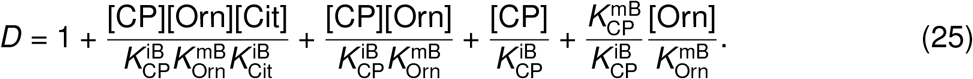

The model included the enzymatic reaction catalyzed by ArcB, CP hydrolysis with velocity 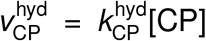,and the dynamic interconversion between P_i_ and dihydrogen phosphate 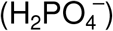. The kinetic parameters of equation (24), with 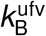set to 89*/*6 s^*−*1^,^46^were obtained by fitting this model to the enzymatic assay data (Supplementary Table 1).

### *L. lactis* carbamate kinase (ArcC1) characterization

Carbamate kinase catalyzes the conversion of CP and MgADP into MgATP and carbamate. Enzymatic assays were performed using CP and MgADP as substrates (Fig. 4d). ArcC1 activity was measured indirectly via ATP luminescence using the ATPlite− Luminescence Assay System. A flat-bottom 96-well white plate (Greiner Bio-One International GmbH) was prepared with 125 µL of diluted mammalian cell lysis solution (2:3 ATPlite− mammalian cell lysis solution : MilliQ). An ATP calibration curve was prepared in duplicate with MgATP concentrations of 0 µM, 31.25 µM, 62.5 µM, 125 µM, 250 µM, 0.5 mM, 1 mM, and 2 mM. When testing CP dependence, MgADP was maintained at 5 mM, and when testing MgADP dependence, CP was maintained at 5 mM. Reaction mixtures contained 5 mM MgCl_2_ in 50 mM KP_i_ buffer at pH 7.0 and were pre-incubated at 30 °C for 1 min to minimize CP degradation. Reactions were initiated by adding ArcC1 to a final concentration of 50 nM. Samples of 5 µL were collected at 0, 10, 20, 40, 60, 80, 100, and 120 s, quenched in the prepared 96-well plate, and diluted with 20 µL of MilliQ water. 50 µL of luciferase solution was added to each well, and the plate was dark-adapted for 10 min in a BioTek Synergy H1MD microplate reader before luminescence was measured with a gain of 135.

*L. lactis* ArcC1 exhibited positive cooperativity between its two nucleotide-binding sites, a feature also observed in another ArcC.^47^ArcC1 activity was therefore modeled using a generalized Hill equation:^48, 49^

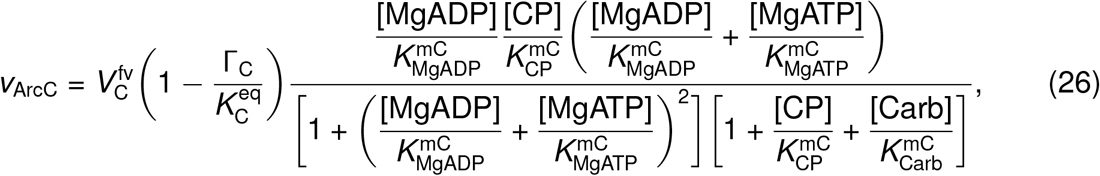

where 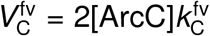 is the limiting velocity in the favorable direction, that is, the direction of MgATP formation. The rate law contains two catalytic rates and four Michaelis constants, constrained by the equilibrium constant 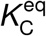through a Haldane relationship.^32^ 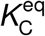was set to 6.0, as obtained at 0.25 M ionic strength and pH 7.0.^50^

Enzyme activity was calculated from the initial phase of the reaction (10 s) and used for curve fitting (Supplementary Table 1). No cooperativity was observed between the two CP binding sites within the concentration ranges explored, despite reports of negative cooperativity.^47^The kinetic parameters of the reverse reaction were inferred by fitting the ADI pathway model to experimental data obtained in vesicles.

### *L. sakei* arginine-ornithine antiporter (ArcD) characterization

*L. sakei* ArcD was characterized in proteoliposomes with external arginine and internal ornithine as substrates (Figs. 4e and 4f and Supplementary Fig. 8). We purified and reconstituted *L. sakei* ArcD at a lipid-to-protein ratio of 400:1 (w/w) in DOPE:DOPG:DOPC (25:25:50 % mol) liposomes as described before.^28^For encapsulation, proteoliposomes resuspended to a lipid concentration of 25 mg mL^*−*1^in 50 mM KP_i_ buffer at pH 7.0, 2 mM DTT, and L-ornithine were subjected to 5 freeze-thaw cycles, with thawing performed in an ice-water bath at 10 °C. Proteoliposomes were then extruded 13× through a 400 nm polycarbonate filter pre-equilibrated with 50 mM KP_i_ buffer at pH 7.0 and 2 mM DTT containing L-ornithine, washed, and collected by centrifugation (20 min, 325 000 ×*g*, 4 °C) in the same buffer without L-ornithine. L-ornithine was encapsulated at concentrations of 0.5, 2.5, 5.0, and 10 mM. Uptake assays were initiated with a 50× dilution of proteoliposomes (200 mg mL^*−*1^ lipid concentration) in pre-heated (30 °C) reaction buffer containing ^14^C-L-arginine, 2 mM DTT, and 50 mM KP_i_ buffer at pH 7.0, with external radiolabeled L-arginine varied from 1 to 100 µM using a ^14^C-L-arginine stock with a specific activity of 325 mCi mmol^*−*1^. Samples of 100 µL were taken at 0, 5, 10, 25, 40, 60, and 90 s, diluted into 2 mL of ice-cold 50 mM KP_i_ buffer at pH 7.0, and filtered through a 0.45 µm cellulose nitrate filter. The filter was washed with 2 mL of ice-cold 50 mM KP_i_ buffer at pH 7.0 and dissolved in Ultima Gold MV scintillation liquid (Perkin Elmer). Radioactivity was quantified in a Tri-Carb 2800TR scintillation counter (PerkinElmer).

*L. sakei* ArcD was assumed to operate via a ping-pong transport mechanism, similar to its *L. lactis* homolog^52^ (Supplementary Fig. 7). ArcD-mediated transport was therefore modeled using the rate law:^53^

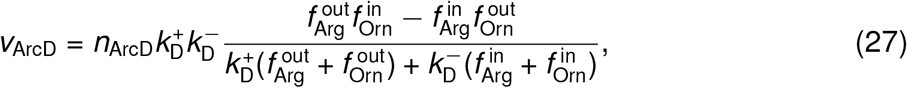

where *n*_ArcD_ is the amount of ArcD in the membrane, and 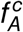 is the fractional concentration of amino acid *A* ∈ {Arg, Orn} in compartment *c* ∈ {out, in}, defined as

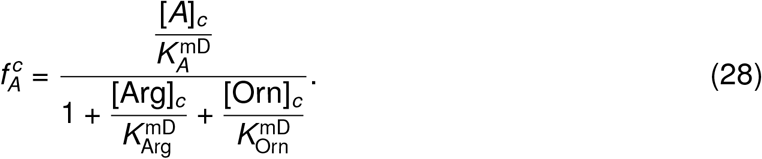

Under the conditions [Arg]_in_ = [Orn]_out_ = 0 and 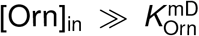, the rate law simplifies to Michaelis-Menten kinetics with catalytic rate 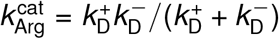, which was set to 6.0 s^*−*1^ based on a previous characterization,^28^ allowing 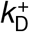 to be calculated from 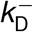.

Each experimental sample had a total volume of 100 µL with a lipid concentration of 3.27 g L^*−*1^. Given a specific internal volume of 2.7 µL per mg lipid,^29^ the internal vesicle volume per sample was calculated as *V*_in_ = 0.8829 µL. The time-dependent uptake of arginine was described by d[Arg]*/*d*t* = *V*_ArcD_*/V*_in_. Fitting this model to the experimental data yielded the kinetic parameters of equation (27) (Supplementary Table 1).

### Mitochondrial ATP-ADP carrier (AAC) modeling

The AAC rate law was derived from literature-reported kinetic parameters rather than direct experimental characterization. The mitochondrial ATP-ADP carrier functions through a ping-pong transport mechanism,^54, 55^ with conformational changes as the rate-limiting steps.^56, 57^ Equal ADP affinities were assumed for the outward- and inward-facing conformations, and the same assumption was applied to ATP. Under these assumptions, AAC-mediated transport was modeled using the rate law:^53^

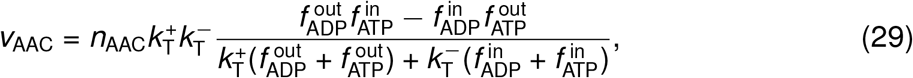

where *n*_AAC_ is the amount of AAC in the membrane, and 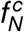 denotes the fractional concentration of nucleotide *N* ∈ {ADP, ATP} in compartment *c* ∈ {out, in}.

Under saturating [ATP]_in_ and with [ADP]_in_ = [ATP]_out_ = 0, equation (29) reduces to Michaelis-Menten kinetics, with Michaelis constant and catalytic rate given by

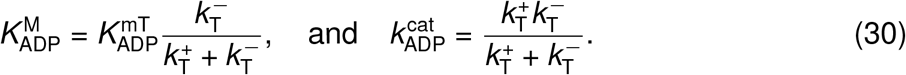

The kinetic parameters 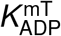 and 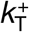 were determined from 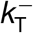 using these relations, with 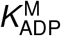 and 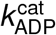 set to 2.2 µM and 1.9 s^*−*1^, respectively.^27^ The ATP affinity and 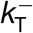 were treated as free parameters to be estimated from vesicle data.

### Reconstitution of the ADI pathway in vesicles

*L. sakei* ArcD and the mitochondrial ATP-ADP carrier from *Thermothelomyces thermophila* (AAC) were co-reconstituted in DOPE:DOPG:DOPC (25:25:50 % mol) liposomes as described before.^25^The proteoliposomes were encapsulated with either 0.5 mM or 10 mM L-ornithine, 10 mM MgCl_2_, 10 mM potassium ADP, 2 mM DTT, 5.8 µM PercevalHR, 1 µM ArcA, 2 µM ArcB, and 5 µM ArcC1 in 50 mM KP_i_ buffer at pH 7.0 by 5 cycles of freeze-thawing. Encapsulated proteoliposomes were extruded 13× through a 400 nm polycarbonate filter and washed 3× by centrifugation at 325 000 ×*g* for 20 min at 4 ^*°*^C and resuspension in 6 mL of external solution containing 50 mM KP_i_ buffer at pH 7.0, 2 mM DTT, and 58 mM NaCl for osmotic balancing. Proteoliposomes were finally resuspended in the same buffer to a lipid concentration of 5.5 mg mL^*−*1^.

### Fluorescence measurements of encapsulated PercevalHR

Proteoliposomes were diluted 2× in external solution, and the excitation spectrum between 400 nm and 520 nm (emission at 550 nm) was recorded at intervals of 1 min. The ADI pathway was initiated by the external addition of 10 mM L-arginine. When indicated, external potassium ADP was added. Valinomycin and nigericin (both diluted in DMSO) were present throughout the experiment at 1 µM. Fluorescence measurements were performed at 30 ^*°*^C in an FP-8350 spectrofluorometer (JASCO, Inc.) with excitation and emission bandwidths of 5 nm.

### PercevalHR calibration curve within liposome

To exclude any external factors, DOPE:DOPG:DOPC (25:25:50 % mol) liposomes were prepared without proteins and encapsulated using the same mixture and method as described above, except that the proteins, other than PercevalHR, were replaced by their storage buffer (50 mM KP_i_ at pH 7.0, 100 mM KCl), and ATP-to-ADP ratios were varied (0.1, 0.4, 1, 4, 10, Fluorescence excitation ratios (F_500_*/*F_430_) were determined by measuring an excitation spectrum between 400 nm and 520 nm (emission at 550 nm) in the presence of 1 µM valinomycin and nigericin (both diluted in DMSO) at 30 ^*°*^C in an FP-8350 spectrofluorometer (JASCO, Inc.) with excitation and emission bandwidths of 5 nm. Experimental data were fitted using a sigmoidal dose-response with non-cooperative binding, yielding a sigmoidal calibration curve (Supplementary Fig. 9):

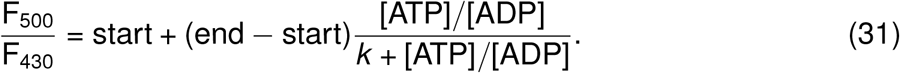

### Relating fluorescence ratio to ATP production rate

At fixed total nucleotide concentration, *A*_tot_ = [ADP] + [ATP], the time derivative of the fluorescence ratio (equation (31)) is related to the ATP production rate by

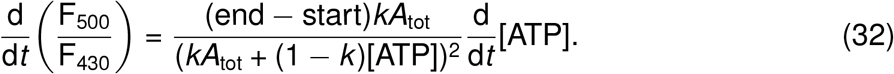

### Development of the kinetic model of the ADI pathway

The ADI pathway was encapsulated within large-unilamellar vesicles (LUVs) with ArcD and AAC reconstituted in the membrane. Each experimental sample contained 0.34 mg total lipids, with a lipid-to-protein ratio of 200:1 (w/w) for the reconstitution of ArcD (total amount of 31.4 pmol) and AAC (51.0 pmol). The total internal vesicle volume in the 122.5 µL reaction volume was *V*_in_ = 0.893 µL,^29^ corresponding to 1.8 · 10^10^ spherical vesicles with an average radius *r*_v_ = 226 nm. For modeling purposes, these vesicles were treated as an equivalent single vesicle with internal volume *V*_in_ and surface area *A*_m_ = 119 cm^2^. Internally, the vesicles contained the ATP/ADP ratio sensor PercevalHR, ornithine, and MgADP in 50 mM KP_i_ buffer at pH 7.0. The reaction was initiated by supplying arginine to the external medium, also 50 mM KP_i_ buffer at pH 7.0. Valinomycin and nigericin were included at 1 µM throughout the experiment.

In addition to the enzyme-catalyzed reactions, several spontaneous processes were included in the model. Ammonium ions produced by ArcA were assumed to rapidly equilibrate with H^+^ and NH_3_ (equations (S.47) and (S.48)), with the latter freely diffusing across the membrane. The H^+^ left behind rapidly equilibrates with the external medium via the ionophores, and the large volume and high buffering capacity of the medium prevent significant pH changes; constant pH was therefore assumed. Interactions among Mg^2+^ and nucleotides were modeled as a dynamic equilibrium (equations (S.51) and (S.52)). Cyanate and carbamate hydrolysis to NH_3_ and 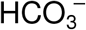 were modeled by simple mass-action kinetics. A list of the considered side reactions and their parameters is provided in Supplementary Table 2.

Ammonia (NH_3_) and carbon dioxide (CO_2_) diffusion across the membrane was described by the Goldman-Hodgkin-Katz flux equation.^58^ The ammonia diffusion rate was expressed as

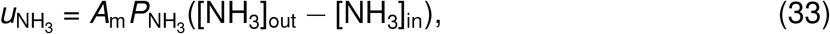

With 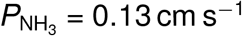 based on measurements in egg PC-decane bilayers.^59^ 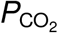 was set to 0.16 cm s^*−*1^ based on measurements on artificial phospholipid vesicles without cholesterol.^60^

Mass-balance equations were written for all intracellular and extracellular species, assembling these components into a unified dynamical model comprising 26 coupled differential equations (equations (S.20)–(S.45)). The remaining parameters were estimated by fitting the model to the measured ATP-to-ADP ratios (Supplementary Fig. 10 and Supplementary Table 3).

### Parameter estimation

Model parameters were estimated by minimizing the residual sum of squares (RSS) between model predictions and experimental data. The Differential Evolution algorithm^61^was used to find the parameter set 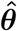 minimizing RSS(***θ***), a general-purpose global optimizer based on empirical evolutionary rules.

Parameter uncertainties were estimated using the likelihood confidence region method,^62^ which defines a confidence region with confidence level 1 − *α* as

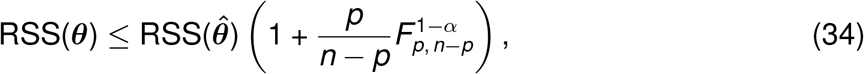

where *p* is the number of parameters, *n* is the number of experimental points, and 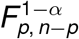 is the *F* −distribution with *p* and *n* − *p* degrees of freedom. The interquartile ranges of this confidence region were used to estimate parameter uncertainties. As an example, the confidence region for the ArcA characterization is shown in Supplementary Fig. 4b.

## Acknowledgements

This work was supported by the Netherlands Organisation for Scientific Research (NWO) through the projects *Building a Synthetic Cell* (BaSyC; grant No. 024.003.019) and *Synthetic non-equilibrium cell-like systems for fuel and pH homeostasis* (grant No. OCENW.M.22.318), as well as by the European Research Council (ERC) through the project *MetaDivide* (grant No. 101167181).

The authors thank Edmund Kunji and Martin King (Medical Research Council Mitochondrial Biology Unit, University of Cambridge) for providing mitochondrial membranes containing AAC from *Thermothelomyces thermophila*.

## Author contributions

MH, BP, YBB, and EPHJ conceived the study and designed the research. YBB developed the coarse-grained models, contributed to the development of the ADI pathway model, performed data analysis, and wrote the manuscript. EPHJ performed enzymatic assays, conducted the PercevalHR calibration and full ADI pathway experiments, analyzed data, and edited the manuscript. MPR contributed to experimental design, performed ArcA and ArcB enzymatic assays, and carried out ArcD reconstitution and characterization. DAJG contributed to the development of the ADI pathway model and the design of vesicle-based experiments. JC contributed to experimental design, protein purification and reconstitution, enzymatic assays, and manuscript editing. MU contributed to the PercevalHR calibration experiments and AAC purification. MH and BP supervised the project, secured funding and resources, and edited the manuscript.

