## Supplemental information for "Accessible Gibbs energy at metabolic activation limits long-term cell growth"

### Supplementary methods

#### Development of the uniport-ATP-coupled cell model

In the uniport-ATP-coupled cell model (Fig. 1c), uniport transport was modeled as a reversible reaction,

$$v_\mathrm{t} = E_\mathrm{t} k_\mathrm{t}^\mathrm{cat} \frac{[\mathrm{S}]_\mathrm{out} - [\mathrm{S}]_\mathrm{in}}{K_\mathrm{t}^\mathrm{mS} + [\mathrm{S}]_\mathrm{out} + [\mathrm{S}]_\mathrm{in}}, \tag{S.1}$$

assuming equal catalytic rate constants for the influx and efflux directions and equal affinity of the binding site for the external and internal substrates. In the cytosol, $[\mathrm{S}]_\mathrm{in}$ is phosphorylated in an ATP-dependent reaction, modeled as

$$v_\mathrm{p} = E_\mathrm{p} k_\mathrm{p}^\mathrm{cat} \frac{[\mathrm{ATP}]}{K_\mathrm{p}^\mathrm{mATP} + [\mathrm{ATP}]} \frac{[\mathrm{S}]_\mathrm{in}}{K_\mathrm{p}^\mathrm{mS} + [\mathrm{S}]_\mathrm{in}}. \tag{S.2}$$

During catabolism, an extra cytosolic $\mathrm{P_i}$ is added to the intermediate I, and the two phosphate groups it now carries are transferred to ADP, yielding two ATP molecules and the precursor P. The catabolic reaction was modeled as

$$v_\mathrm{c} = E_\mathrm{c} k_\mathrm{c}^\mathrm{cat} \left( \frac{[\mathrm{ADP}]}{K_\mathrm{c}^\mathrm{mADP} + [\mathrm{ADP}]} \right)^2 \frac{[\mathrm{I}]}{K_\mathrm{c}^\mathrm{mI} + [\mathrm{I}]}. \tag{S.3}$$

Anabolism was modeled as

$$v_\mathrm{a} = E_\mathrm{a} k_\mathrm{a}^\mathrm{cat} \frac{[\mathrm{ATP}]}{K_\mathrm{a}^\mathrm{mATP} + [\mathrm{ATP}]} \frac{[\mathrm{P}]}{K_\mathrm{a}^\mathrm{mP} + [\mathrm{P}]}. \tag{S.4}$$

Following the approach described for the antiport-coupled model, the total enzyme concentration was obtained as

$$\frac{E_\mathrm{tot}}{J} = \mathcal{A} + \frac{1}{k_\mathrm{t}^\mathrm{cat}} \frac{K_\mathrm{t}^\mathrm{mS} + 2[\mathrm{S}]_\mathrm{in}}{[\mathrm{S}]_\mathrm{out} - [\mathrm{S}]_\mathrm{in}} + \frac{\mathcal{B}_\mathrm{p}^\mathrm{S}}{[\mathrm{S}]_\mathrm{in}} + \frac{\mathcal{B}_\mathrm{p}^\mathrm{ATP}}{[\mathrm{ATP}]} + \frac{\mathcal{B}_\mathrm{p}^\mathrm{S} \mathcal{B}_\mathrm{p}^\mathrm{ATP} k_\mathrm{p}^\mathrm{cat}}{[\mathrm{S}]_\mathrm{in}[\mathrm{ATP}]} + \frac{\mathcal{B}_\mathrm{c}^\mathrm{I}}{[\mathrm{I}]} + $$

$$+ \frac{\mathcal{B}_\mathrm{c}^\mathrm{ADP}}{Q_0 - [\mathrm{ATP}]} \left( 1 + \frac{K_\mathrm{c}^\mathrm{mI}}{[\mathrm{I}]} \right) \left( 2 + \frac{K_\mathrm{c}^\mathrm{mADP}}{Q_0 - [\mathrm{ATP}]} \right) + \frac{\mathcal{B}_\mathrm{a}^\mathrm{P}}{[\mathrm{P}]} + \frac{\mathcal{B}_\mathrm{a}^\mathrm{ATP}}{[\mathrm{ATP}]} + \frac{\mathcal{B}_\mathrm{a}^\mathrm{P} \mathcal{B}_\mathrm{a}^\mathrm{ATP} k_\mathrm{a}^\mathrm{cat}}{[\mathrm{P}][\mathrm{ATP}]}. \tag{S.5}$$

A constraint on the total pool of internal metabolites was enforced by defining a Lagrangian function with Lagrange multiplier $\lambda$ as

$$\mathcal{L}_\lambda = E_\mathrm{tot} - \lambda\big([\mathrm{S}]_\mathrm{tot} - Q_0 - [\mathrm{S}]_\mathrm{in} - [\mathrm{I}] - [\mathrm{P}]\big). \tag{S.6}$$

Minimizing $\mathcal{L}_\lambda$ with respect to $[\mathrm{S}]_\mathrm{in}$, [I], and [P] yielded

$$\frac{1}{x^2} = \left( \frac{\mathcal{N}_1}{[\mathrm{S}]_\mathrm{in}} \right)^2 - \frac{1/k_\mathrm{t}^\mathrm{cat}}{[\mathrm{S}]_\mathrm{out} - [\mathrm{S}]_\mathrm{in}} \left( 2 + \frac{K_\mathrm{t}^\mathrm{mS} + 2[\mathrm{S}]_\mathrm{in}}{[\mathrm{S}]_\mathrm{out} - [\mathrm{S}]_\mathrm{in}} \right), \quad [\mathrm{I}] = x\mathcal{N}_2, \quad \text{and} \quad [\mathrm{P}] = x\mathcal{N}_3, \tag{S.7}$$

where $x = \sqrt{J/\lambda}$ and the numerators $\mathcal{N}_i$ are defined as

$$\mathcal{N}_1 = \sqrt{\mathcal{B}_{\rm p}^{\rm S}\left(1 + \frac{K_{\rm p}^{\rm mATP}}{[\rm ATP]}\right)}, \tag{S.8}$$

$$\mathcal{N}_2 = \sqrt{\mathcal{B}_{\rm c}^{\rm I}\left[1 + \frac{K_{\rm c}^{\rm mADP}}{[\rm ADP]}\left(2 + \frac{K_{\rm c}^{\rm mADP}}{[\rm ADP]}\right)\right]}, \quad \text{and} \quad \mathcal{N}_3 = \sqrt{\mathcal{B}_{\rm a}^{\rm P}\left(1 + \frac{K_{\rm a}^{\rm mATP}}{[\rm ATP]}\right)}. \tag{S.9}$$

Minimizing $\mathcal{L}_\lambda$ with respect to [ATP] yielded

$$\gamma = 2\left(\frac{[\rm ATP]}{[\rm ADP]}\right)^2\left(1 + \frac{K_{\rm c}^{\rm mADP}}{[\rm ADP]}\right), \tag{S.10}$$

where $\gamma$ is defined as

$$\gamma = \left[\frac{\mathcal{B}_{\rm p}^{\rm ATP}}{\mathcal{B}_{\rm c}^{\rm ADP}}\left(1 + \frac{K_{\rm p}^{\rm mS}}{[\rm S]_{\rm in}}\right) + \frac{\mathcal{B}_{\rm a}^{\rm ATP}}{\mathcal{B}_{\rm c}^{\rm ADP}}\left(1 + \frac{K_{\rm a}^{\rm mP}}{[\rm P]}\right)\right]\left(1 + \frac{K_{\rm c}^{\rm mI}}{[\rm I]}\right)^{-1}. \tag{S.11}$$

Equations S.7 and S.10 form a system of transcendental equations that was solved numerically to obtain the steady-state metabolite concentrations minimizing $E_{\rm tot}$. These concentrations were substituted into equation (S.5) to compute the normalized growth rate ($\mu \propto J/E_{\rm tot}$) as a function of the external substrate concentration and $Q_0$.

For the simulations (Fig. 1f and Supplementary Fig. 1), we used the same catalytic rate constants as in the antiport-coupled model, and the Michaelis constants $K_{\rm t}^{\rm mS} = 50\,\mu$M, $K_{\rm p}^{\rm mATP} = K_{\rm p}^{\rm mS} = K_{\rm a}^{\rm mATP} = 0.1$ mM, and $K_{\rm c}^{\rm mADP} = K_{\rm c}^{\rm mI} = K_{\rm a}^{\rm mP} = 0.5$ mM. Additionally, we set $[\rm S]_{\rm tot} = 100$ mM, and considered $[\rm ADP]_0 = [\rm S]_{\rm in}^0 = 1$ pM, and $[\rm P_i] = 10$ mM.

#### Development of the PTS-coupled cell model

In the PTS-coupled cell model (Fig. 1d), catabolism and anabolism were described using the same rate laws as in the uniport-ATP-coupled model, while substrate uptake and phosphorylation were combined into a single ATP-driven reaction, modeled as

$$v_{\rm t} = E_{\rm t} k_{\rm t}^{\rm cat} \frac{[\rm ATP]}{K_{\rm t}^{\rm mATP} + [\rm ATP]} \frac{[\rm S]_{\rm out}}{K_{\rm t}^{\rm mS} + [\rm S]_{\rm out}}. \tag{S.12}$$

The Lagrangian was defined as $\mathcal{L}_\lambda = E_{\rm tot} - \lambda([\rm S]_{\rm tot} - Q_0 - [\rm I] - [\rm P])$, with the total enzyme concentration given by

$$\begin{aligned}\frac{E_{\rm tot}}{J} = \mathcal{A} &+ \frac{\mathcal{B}_{\rm t}^{\rm S}}{[\rm S]_{\rm out}} + \frac{\mathcal{B}_{\rm t}^{\rm ATP}}{[\rm ATP]} + \frac{\mathcal{B}_{\rm t}^{\rm S}\mathcal{B}_{\rm t}^{\rm ATP}k_{\rm t}^{\rm cat}}{[\rm S]_{\rm out}[\rm ATP]} + \frac{\mathcal{B}_{\rm c}^{\rm I}}{[\rm I]} + \\ &+ \frac{\mathcal{B}_{\rm c}^{\rm ADP}}{Q_0 - [\rm ATP]}\left(1 + \frac{K_{\rm c}^{\rm mI}}{[\rm I]}\right)\left(2 + \frac{K_{\rm c}^{\rm mADP}}{Q_0 - [\rm ATP]}\right) + \frac{\mathcal{B}_{\rm a}^{\rm P}}{[\rm P]} + \frac{\mathcal{B}_{\rm a}^{\rm ATP}}{[\rm ATP]} + \frac{\mathcal{B}_{\rm a}^{\rm P}\mathcal{B}_{\rm a}^{\rm ATP}k_{\rm a}^{\rm cat}}{[\rm P][\rm ATP]}.\end{aligned} \tag{S.13}$$

Minimizing $\mathcal{L}_\lambda$ with respect to [I] and [P] yielded

$$[\mathrm{I}] = \frac{\mathcal{N}_2}{\mathcal{N}_2 + \mathcal{N}_3}([\mathrm{S}]_{\mathrm{tot}} - Q_0), \quad \text{and} \quad [\mathrm{P}] = \frac{\mathcal{N}_3}{\mathcal{N}_2 + \mathcal{N}_3}([\mathrm{S}]_{\mathrm{tot}} - Q_0), \tag{S.14}$$

where $\mathcal{N}_2$ and $\mathcal{N}_3$ are defined in equation (S.9). Minimizing $\mathcal{L}_\lambda$ with respect to [ATP] yielded an equation of the same form as equation (S.10), but with $\gamma$ defined as

$$\gamma = \left[\frac{\mathcal{B}_{\mathrm{t}}^{\mathrm{ATP}}}{\mathcal{B}_{\mathrm{c}}^{\mathrm{ADP}}}\left(1 + \frac{K_{\mathrm{t}}^{\mathrm{mS}}}{[\mathrm{S}]_{\mathrm{out}}}\right) + \frac{\mathcal{B}_{\mathrm{a}}^{\mathrm{ATP}}}{\mathcal{B}_{\mathrm{c}}^{\mathrm{ADP}}}\left(1 + \frac{K_{\mathrm{a}}^{\mathrm{mP}}}{[\mathrm{P}]}\right)\right]\left(1 + \frac{K_{\mathrm{c}}^{\mathrm{mI}}}{[\mathrm{I}]}\right)^{-1}. \tag{S.15}$$

For the simulations (Fig. 1g and Supplementary Fig. 2), we used the catalytic rate constants $k_{\mathrm{t}}^{\mathrm{cat}} = 5/6\,\mathrm{s}^{-1}$, $k_{\mathrm{c}}^{\mathrm{cat}} = 5\,\mathrm{s}^{-1}$, and $k_{\mathrm{a}}^{\mathrm{cat}} = 0.1\,\mathrm{s}^{-1}$, and the Michaelis constants $K_{\mathrm{t}}^{\mathrm{mS}} = 50\,\mu\mathrm{M}$, $K_{\mathrm{t}}^{\mathrm{mATP}} = K_{\mathrm{a}}^{\mathrm{mATP}} = 0.1\,\mathrm{mM}$, and $K_{\mathrm{c}}^{\mathrm{mADP}} = K_{\mathrm{c}}^{\mathrm{mI}} = K_{\mathrm{a}}^{\mathrm{mP}} = 0.5\,\mathrm{mM}$. Additionally, we set $[\mathrm{S}]_{\mathrm{tot}} = 100\,\mathrm{mM}$, and considered $[\mathrm{ADP}]_0 = 1\,\mathrm{pM}$.

#### Quantifying accessible Gibbs energy of the ATP-coupled models

In the uniport-ATP-coupled model, Gibbs energy is stored in two distinct forms. First, concentration differences across the membrane store Gibbs energy that drives substrate uptake through uniport transport. Second, Gibbs energy is stored in high ATP-to-ADP ratios and is dissipated via ATP hydrolysis. Both contributions depend on the system state and were therefore included in the computation of the accessible Gibbs energy. In the PTS-coupled model, substrate uptake is directly coupled to ATP-dependent phosphorylation, and the accessible Gibbs energy was determined solely by the Gibbs energy associated with ATP hydrolysis. In both models, each contribution was quantified using the same procedure as for the antiport-coupled model.

In the uniport-ATP-coupled model, the Gibbs energy change per mole of transported substrate is

$$\Delta G_{\mathrm{uniport}} = RT \log\left(\frac{[\mathrm{S}]_{\mathrm{in}}}{[\mathrm{S}]_{\mathrm{out}}}\right). \tag{S.16}$$

Following the same procedure as for the antiport model, we considered the transport of an amount of substrate that changes the intracellular concentration according to $[\mathrm{S}]_{\mathrm{in}} = [\mathrm{S}]_{\mathrm{in}}^* + \xi$. Integrating $d\mathcal{E} = -\Delta G_{\mathrm{uniport}}\, d\xi$ up to the maximum concentration change $\xi_{\max}$ gave the accessible Gibbs energy of uniport:

$$\frac{\mathcal{E}}{RT} = \xi_{\max}\left[1 + \log\left(\frac{[\mathrm{S}]_{\mathrm{out}}}{[\mathrm{S}]_{\mathrm{in}}^* + \xi_{\max}}\right)\right] + [\mathrm{S}]_{\mathrm{in}}^* \log\left(\frac{[\mathrm{S}]_{\mathrm{in}}^*}{[\mathrm{S}]_{\mathrm{in}}^* + \xi_{\max}}\right). \tag{S.17}$$

The maximum concentration change $\xi_{\max} \geq 0$ is constrained by three independent upper bounds. The first bound is obtained by integrating $d\mathcal{E}$ up to the point at which $\Delta G_{\mathrm{uniport}}$ vanishes, corresponding to complete dissipation of the transmembrane chemical gradient:

$\xi_{max}$ = $[S]_{out} - [S]^*_{in}$. The second bound arises from the finite total pool of internal metabolites: $\xi_{max} = [S]_{tot} - Q_0 - [S]^*_{in}$. The third bound follows from ATP availability: although uniport transport itself does not consume ATP, only substrate that can be phosphorylated contributes to the accessible Gibbs energy, imposing $\xi_{max} \leq [ATP]^* - [S]^*_{in}$. In practice, $\xi_{max}$ was set for each condition by the most restrictive of these three bounds.

For ATP hydrolysis, the Gibbs energy change per mole of ATP hydrolyzed is

$$\Delta G_{hydrolysis} = \Delta G^{\ominus}_{app} + RT \log\left(\frac{[ADP]}{[ATP]}\right), \tag{S.18}$$

where $\Delta G^{\ominus}_{app}/RT = -12.1 + \log [P_i]$ is the apparent standard Gibbs energy change.[1] In our models, $[P_i]$ was fixed at 10 mM. Following the same procedure as for the antiport-coupled model, the accessible Gibbs energy stored in the ATP/ADP metabolite configuration was obtained as

$$\frac{\mathcal{E}}{RT} = Q_0 \log\left(\frac{1 + K^a_{app}}{Q_0}\right) + [ATP]^* \log\left(\frac{[ATP]^*}{K^a_{app}}\right) + [ADP]^* \log [ADP]^*, \tag{S.19}$$

where $K^a_{app} = \exp(\Delta G^{\ominus}_{app}/RT)$ is the apparent affinity constant for ATP hydrolysis, and we used that $[ADP]^* + [ATP]^* = Q_0$.

#### Samples collection, amino acid derivatization and RP-HPLC analysis

Samples of 60 µL from the enzymatic reactions were taken at discrete time points and transferred to 20 µL of stop solution containing 7% perchloric acid plus 4.5 mM EDTA. Excess acid was neutralized by the addition of 15 µL of neutralization solution containing 1 M KOH and 1 M $KHCO_3$. Samples were frozen and stored at −20 °C overnight or until analysis via HPLC.

Samples were centrifuged in a tabletop centrifuge at 10 000 ×*g* for 5 min at room temperature, and 60 µL of the supernatant was mixed with 87.5 µL of 1 M sodium borate at pH 9.0, 37.5 µL of methanol, and 1.5 µL of diethyl ethoxymethylenemalonate (DEEMM). The derivatization reaction was run for 30 min in a water bath sonicator, followed by incubation at 70 °C for 2 h to degrade excess DEEMM.[2] Samples were cleared by centrifugation at 10 000 ×*g* for 10 min at room temperature, and 100 µL of the supernatant was transferred to a glass vial for RP-HPLC analysis, with absorbance monitored at a wavelength of 280 nm.

Amino acid samples were analyzed by RP-HPLC on a 1260 LC HPLC system (Agilent) composed of a G1311B binary pump, G1329B autosampler, G1316A thermostated column compartment, and a G1315C diode array detector, using a Shimadzu XR-ODS 3 × 75 mm C18 column. Samples were run in a binary gradient between eluent A (25 mM acetate buffer at pH 5.8, 0.02% (w/v) sodium azide) and eluent B (acetonitrile : methanol 8:2 (v/v)) with a flow rate of 0.9 mL min$^{-1}$, at a column temperature of 40 °C and an injection volume of 10 µL. The gradient protocol was run for 20 min as follows: 94% eluent A, 6% eluent B for 2 min;

87% eluent A, 13% eluent B at 2 min; 83% eluent A, 17% eluent B at 8 min; 71% eluent A, 29% eluent B at 9 min; 67% eluent A, 33% eluent B at 12 min; 40% eluent A, 60% eluent B at 12.1 min; and 94% eluent A, 6% eluent B at 16 min. Concentrations of L-ornithine and L-citrulline were determined by interpolation of the corresponding peak area in a calibration curve run with amino acid standards under the same conditions.

#### Development of the kinetic model of the ADI pathway

The system of 26 mass-balance equations describing the dynamics of the encapsulated ADI pathway is:

$$\frac{\mathrm{d}}{\mathrm{d}t}[\mathrm{ADP}]_{\mathrm{out}} = -\frac{v_{\mathrm{AAC}}}{V_{\mathrm{out}}}, \tag{S.20}$$

$$\frac{\mathrm{d}}{\mathrm{d}t}[\mathrm{ATP}]_{\mathrm{out}} = \frac{v_{\mathrm{AAC}}}{V_{\mathrm{out}}}, \tag{S.21}$$

$$\frac{\mathrm{d}}{\mathrm{d}t}[\mathrm{Arg}]_{\mathrm{out}} = -\frac{v_{\mathrm{ArcD}}}{V_{\mathrm{out}}}, \tag{S.22}$$

$$\frac{\mathrm{d}}{\mathrm{d}t}[\mathrm{Carb}]_{\mathrm{out}} = -v_{\mathrm{Carb}}^{\mathrm{out}}, \tag{S.23}$$

$$\frac{\mathrm{d}}{\mathrm{d}t}[\mathrm{CO_2}]_{\mathrm{out}} = -\frac{u_{\mathrm{CO_2}}}{V_{\mathrm{out}}} - v_{\mathrm{CO_2}}^{\mathrm{out}}, \tag{S.24}$$

$$\frac{\mathrm{d}}{\mathrm{d}t}[\mathrm{HCO_3^-}]_{\mathrm{out}} = v_{\mathrm{CO_2}}^{\mathrm{out}} + v_{\mathrm{Carb}}^{\mathrm{out}}, \tag{S.25}$$

$$\frac{\mathrm{d}}{\mathrm{d}t}[\mathrm{NH_3}]_{\mathrm{out}} = -\frac{u_{\mathrm{NH_3}}}{V_{\mathrm{out}}} + v_{\mathrm{NH_4^+}}^{\mathrm{out}} + v_{\mathrm{Carb}}^{\mathrm{out}}, \tag{S.26}$$

$$\frac{\mathrm{d}}{\mathrm{d}t}[\mathrm{NH_4^+}]_{\mathrm{out}} = -v_{\mathrm{NH_4^+}}^{\mathrm{out}}, \tag{S.27}$$

$$\frac{\mathrm{d}}{\mathrm{d}t}[\mathrm{Orn}]_{\mathrm{out}} = \frac{v_{\mathrm{ArcD}}}{V_{\mathrm{out}}}, \tag{S.28}$$

$$\frac{\mathrm{d}}{\mathrm{d}t}[\mathrm{ADP}]_{\mathrm{in}} = \frac{v_{\mathrm{AAC}}}{V_{\mathrm{in}}} + v_{\mathrm{MgADP}}, \tag{S.29}$$

$$\frac{\mathrm{d}}{\mathrm{d}t}[\mathrm{ATP}]_{\mathrm{in}} = -\frac{v_{\mathrm{AAC}}}{V_{\mathrm{in}}} + v_{\mathrm{MgATP}}, \tag{S.30}$$

$$\frac{\mathrm{d}}{\mathrm{d}t}[\mathrm{Arg}]_{\mathrm{in}} = \frac{v_{\mathrm{ArcD}}}{V_{\mathrm{in}}} - v_{\mathrm{ArcA}}, \tag{S.31}$$

$$\frac{\mathrm{d}}{\mathrm{d}t}[\mathrm{Carb}]_{\mathrm{in}} = v_{\mathrm{ArcC}} - v_{\mathrm{Carb}}^{\mathrm{in}} + \tfrac{1}{2}v_{\mathrm{CP}}^{\mathrm{hyd}}, \tag{S.32}$$

$$\frac{\mathrm{d}}{\mathrm{d}t}[\mathrm{Cit}]_{\mathrm{in}} = v_{\mathrm{ArcA}} - v_{\mathrm{ArcB}}, \tag{S.33}$$

$$\frac{d}{dt}[CO_2]_{in} = \frac{u_{CO_2}}{V_{in}} - v^{in}_{CO_2}, \tag{S.34}$$

$$\frac{d}{dt}[CP]_{in} = v_{ArcB} - v_{ArcC} - v^{hyd}_{CP}, \tag{S.35}$$

$$\frac{d}{dt}[HCO_3^-]_{in} = v^{in}_{CO_2} + v^{in}_{Carb} + v_{OCN^-}, \tag{S.36}$$

$$\frac{d}{dt}[H_2PO_4^-]_{in} = -v_{buff}, \tag{S.37}$$

$$\frac{d}{dt}[P_i]_{in} = -v_{ArcB} + v_{buff} + v^{hyd}_{CP}, \tag{S.38}$$

$$\frac{d}{dt}[Mg^{2+}]_{in} = v_{MgADP} + v_{MgATP}, \tag{S.39}$$

$$\frac{d}{dt}[MgADP]_{in} = -v_{ArcC} - v_{MgADP}, \tag{S.40}$$

$$\frac{d}{dt}[MgATP]_{in} = v_{ArcC} - v_{MgATP}, \tag{S.41}$$

$$\frac{d}{dt}[NH_3]_{in} = \frac{u_{NH_3}}{V_{in}} + v^{in}_{NH_4^+} + v^{in}_{Carb} + v_{OCN^-}, \tag{S.42}$$

$$\frac{d}{dt}[NH_4^+]_{in} = v_{ArcA} - v^{in}_{NH_4^+}, \tag{S.43}$$

$$\frac{d}{dt}[OCN^-]_{in} = -v_{OCN^-} + \frac{1}{2}v^{hyd}_{CP}, \tag{S.44}$$

$$\frac{d}{dt}[Orn]_{in} = -\frac{v_{ArcD}}{V_{in}} + v_{ArcB}. \tag{S.45}$$

The velocity of $H_2PO_4^-$ deprotonation was obtained from the fact that the buffer reaction is much faster than the enzyme-catalyzed reactions, maintaining a constant equilibrium state. Assuming a constant pH, the expression for this velocity is:

$$v_{buff} = \frac{v_{ArcB} - v^{hyd}_{CP}}{1 + 10^{pH - pK^a_{buff}}}. \tag{S.46}$$

We derived this expression by substituting Eqs. S.37 and S.38 into the equation obtained by differentiating both sides of $K^a_{buff} = [P_i][H^+]/[H_2PO_4^-]$ with respect to $t$. This approach was also applied to the deprotonation reaction of $NH_4^+$ and $CO_2$ hydrolysis, resulting in

$$v^{out}_{NH_4^+} = \frac{u_{NH_3}/V_{out} - v^{out}_{Carb}}{1 + 10^{pH - pK^a_{NH_4^+}}}, \tag{S.47}$$

$$v^{in}_{NH_4^+} = v_{ArcA} - \frac{v_{ArcA} + u_{NH_3}/V_{in} + v^{in}_{Carb} + v_{OCN^-}}{1 + 10^{pH - pK^a_{NH_4^+}}}, \tag{S.48}$$

$$v_{CO_2}^{out} = -u_{CO_2}/V_{out} - \frac{v_{Carb}^{out} - u_{CO_2}/V_{out}}{1 + 10^{pH - pK_{CO_2}^{a}}}, \tag{S.49}$$

$$v_{CO_2}^{in} = u_{CO_2}/V_{in} - \frac{v_{Carb}^{in} + v_{OCN^-} + u_{CO_2}/V_{in}}{1 + 10^{pH - pK_{CO_2}^{a}}}. \tag{S.50}$$

The interactions among $Mg^{2+}$ and nucleotides establish a dynamic equilibrium. The velocities governing the time evolution of this equilibrium were determined by solving the following system of algebraic equations:

$$\left(K_{MgADP}^{d} + [Mg^{2+}]_{in} + [ADP]_{in}\right) v_{MgADP} + [ADP]_{in} v_{MgATP} = -K_{MgADP}^{d} v_{ArcC} - [Mg^{2+}]_{in} \frac{v_{AAC}}{V_{in}}, \tag{S.51}$$

$$[ATP]_{in} v_{MgADP} + \left(K_{MgATP}^{d} + [Mg^{2+}]_{in} + [ATP]_{in}\right) v_{MgATP} = K_{MgATP}^{d} v_{ArcC} + [Mg^{2+}]_{in} \frac{v_{AAC}}{V_{in}}. \tag{S.52}$$

### Supplementary figures

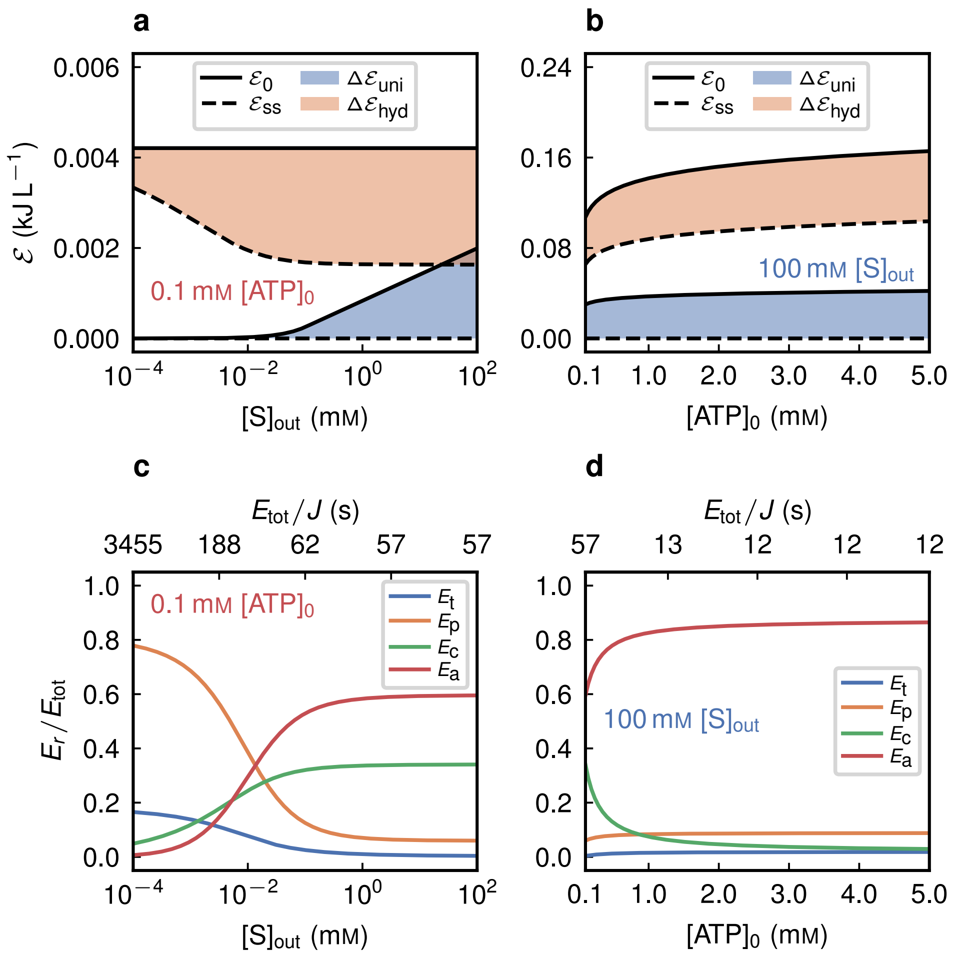


**Supplementary Figure 1: Accessible Gibbs energy limits metabolic activation in the uniport-ATP model.** **a**, Initial accessible Gibbs energy ($\mathcal{E}_0$, calculated from equation (S.17) for uniport, and equation (S.19) for ATP hydrolysis), steady-state accessible Gibbs energy ($\mathcal{E}_{ss}$), and dissipated energy $\Delta\mathcal{E} = \mathcal{E}_0 - \mathcal{E}_{ss}$ as a function of the external substrate concentration $[S]_{out}$, at a limiting initial ATP concentration $[ATP]_0$ = 0.1 mM. **c**, Proteome allocation to transport ($E_t$), phosphorylation ($E_p$), catabolic ($E_c$), and anabolic ($E_a$) enzymes, expressed as fractions of the total enzyme concentration $E_{tot}$, for the simulations in panel **a**. The top axis is annotated with the corresponding total enzyme concentration per unit flux. **b** and **d**, Same analyses as in panels **a** and **c**, respectively, but as a function of $[ATP]_0$ at saturating external substrate concentration $[S]_{out}$ = 100 mM.

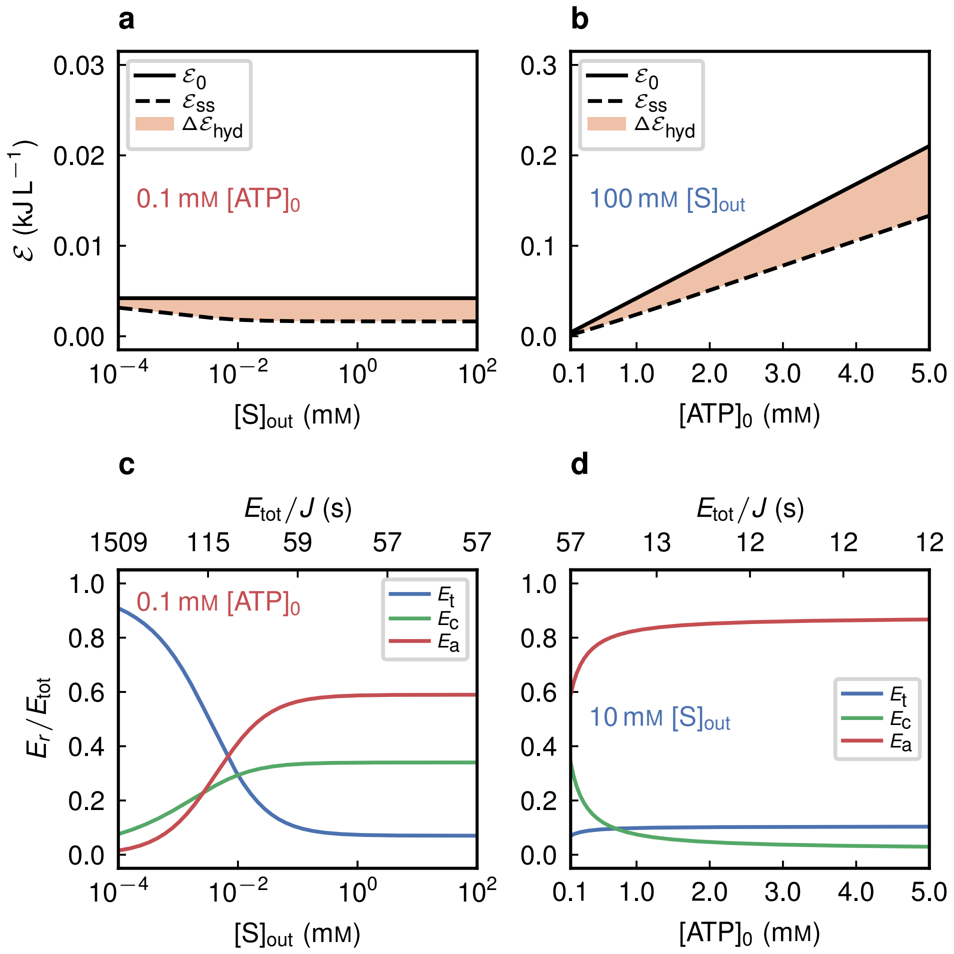


**Supplementary Figure 2: Accessible Gibbs energy limits metabolic activation in the PTS model. a**, Initial accessible Gibbs energy ($\mathcal{E}_0$, calculated from equation (S.19)), steady-state accessible Gibbs energy ($\mathcal{E}_{ss}$), and dissipated energy $\Delta\mathcal{E} = \mathcal{E}_0 - \mathcal{E}_{ss}$ as a function of the external substrate concentration $[S]_{out}$, at a limiting initial ATP concentration $[ATP]_0$ = 0.1 mM. **c**, Proteome allocation to transport ($E_t$), catabolic ($E_c$), and anabolic ($E_a$) enzymes, expressed as fractions of the total enzyme concentration $E_{tot}$, for the simulations in panel **a**. The top axis is annotated with the corresponding total enzyme concentration per unit flux. **b** and **d**, Same analyses as in panels **a** and **c**, respectively, but as a function of $[ATP]_0$ at saturating external substrate concentration $[S]_{out}$ = 100 mM.

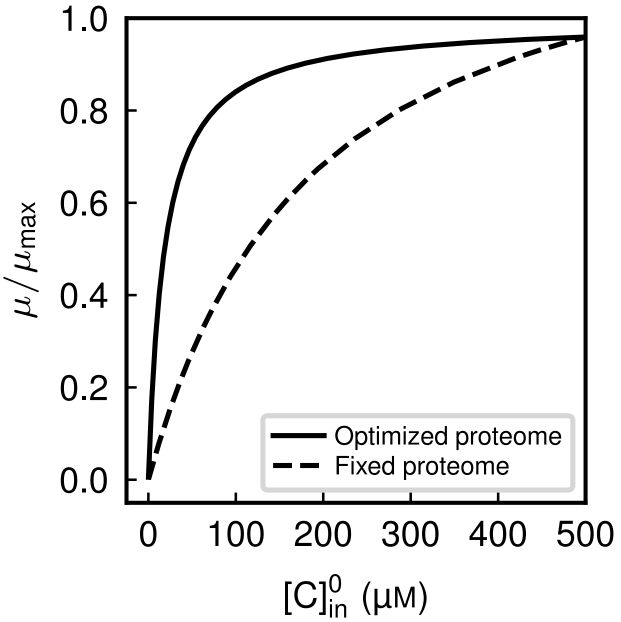


**Supplementary Figure 3: Growth limitation under fixed proteome allocation.** Normalized growth rate as a function of initial countersubstrate concentration, $[C]^0_{in}$, in the antiport-coupled model at saturating external substrate ($[S]_{out}$ = 100 mM). The solid curve shows growth under full proteome optimization, where enzyme allocation and metabolite pools are jointly adjusted at each $[C]^0_{in}$. The dashed curve shows a scenario in which enzyme levels were first determined at high accessible Gibbs energy ($[C]^0_{in}$ = 500 µM) and then held fixed, while the maximal steady-state flux was computed through metabolite reorganization alone.

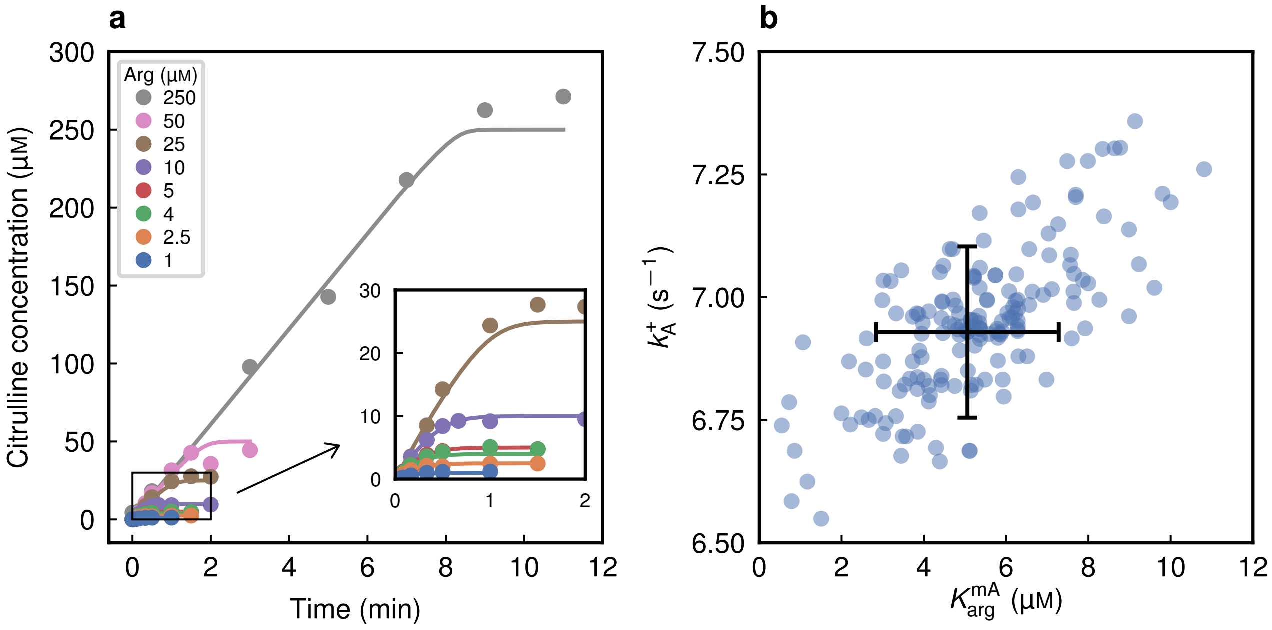


**Supplementary Figure 4: Arginine deiminase characterization.** **a**, Dynamic citrulline formation for initial arginine concentrations ranging from 1 to 250 μM. The model fitted to the experimental data is shown by the solid lines. **b**, Confidence region of the parameters, with a confidence level of $1 - \alpha = 0.95$ (see "Parameter estimation" in the Methods section of the main text).

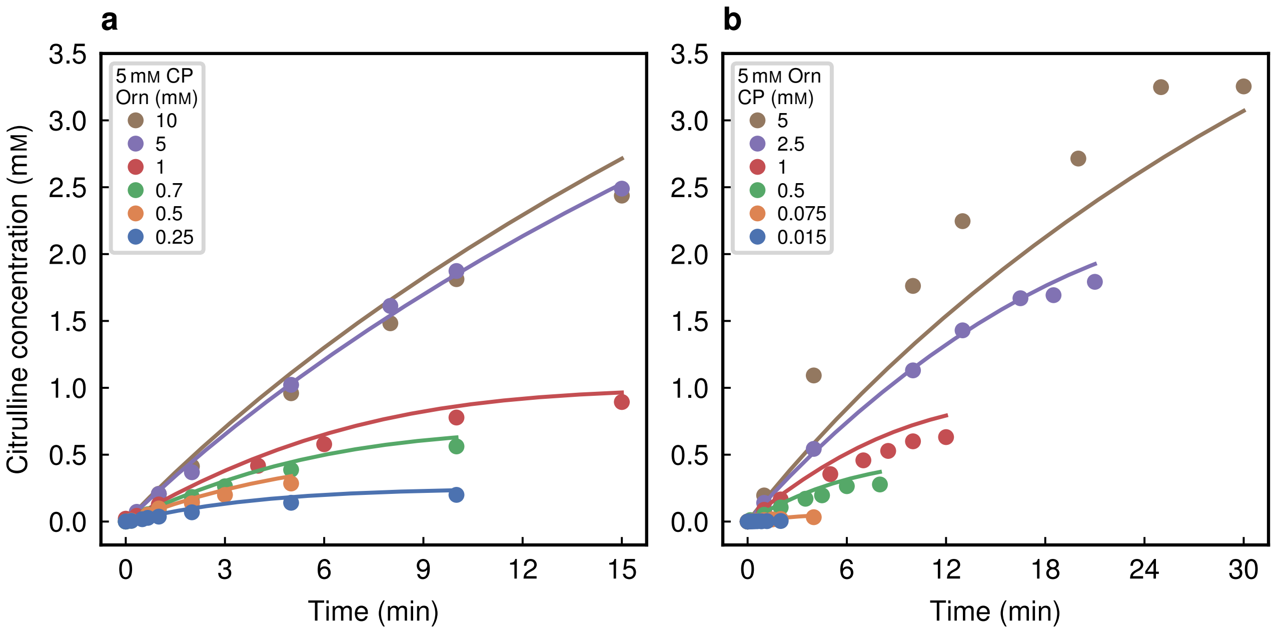


**Supplementary Figure 5: Ornithine transcarbamoylase characterization in the thermodynamically favorable direction.** **a**, Dynamic citrulline formation for initial ornithine concentrations ranging from 0.25 to 10 mM and initial CP concentration of 5 mM. **b**, Dynamic citrulline formation for initial CP concentrations ranging from 0.015 to 5 mM and initial ornithine concentration of 5 mM.

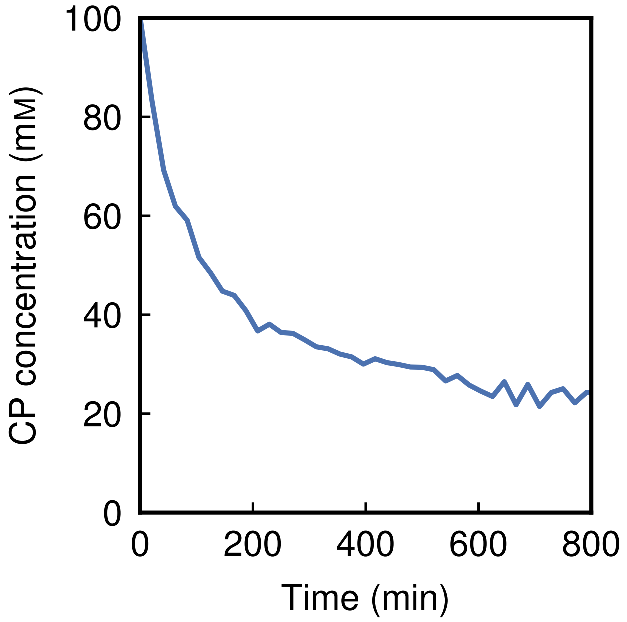


**Supplementary Figure 6: Carbamoyl phosphate degradation characterization.** Dynamic degradation of 100 mM carbamoyl phosphate monitored using $^{31}$P-NMR.

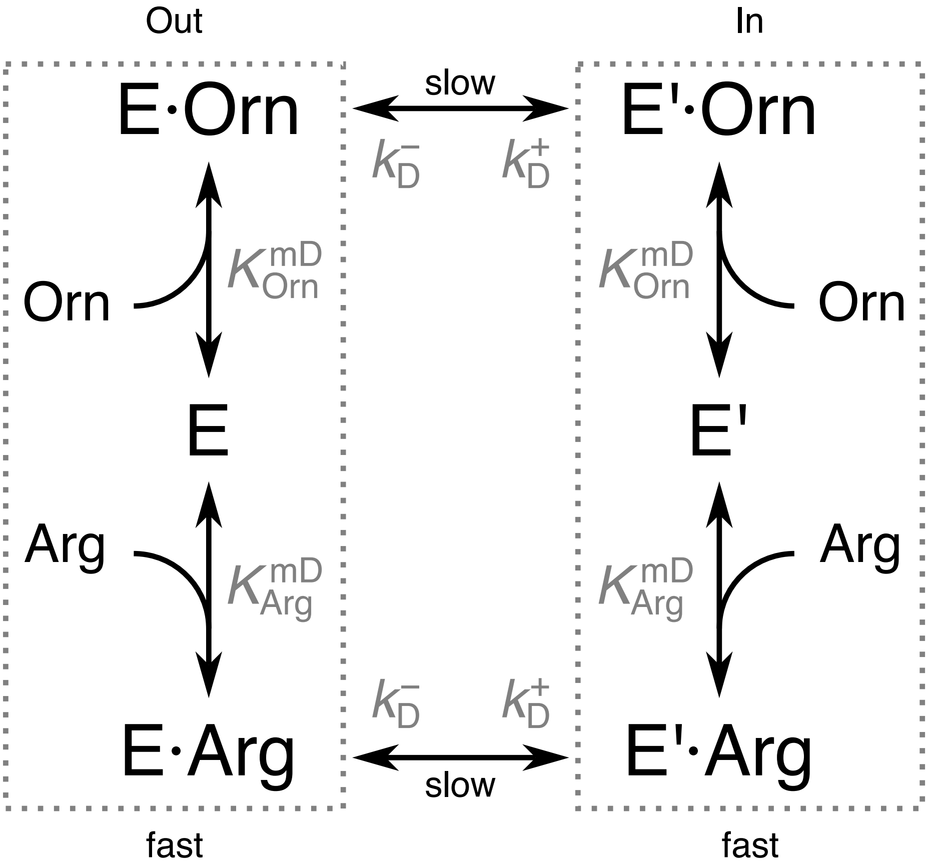


**Supplementary Figure 7: Kinetic diagram of the transport cycle of the arginine-ornithine antiporter.** The state E represents the free carrier, whereas the states E·Arg and E·Orn correspond to the carrier-arginine and carrier-ornithine complexes, respectively. Unprimed states denote the outward-facing conformations of the carrier, while primed states represent the inward-facing conformations. In this model, we assume that the arginine affinities for the outward- and inward-facing conformations are identical, and similarly for ornithine. This assumption simplifies the model significantly without compromising the quality of the fit (not shown). The conformational changes are the rate-limiting steps.

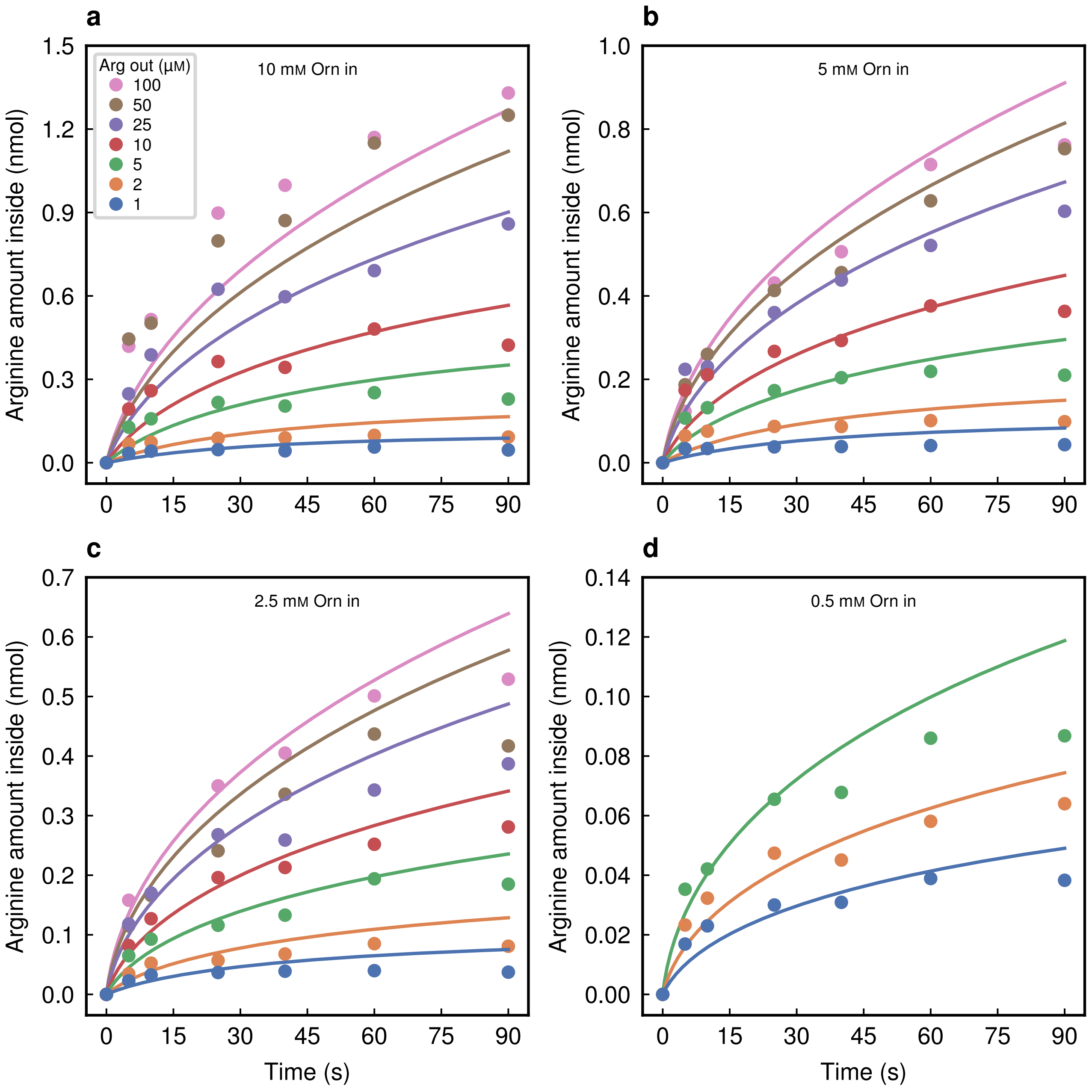


**Supplementary Figure 8: Arginine-ornithine antiporter characterization.** Dynamic arginine uptake in proteoliposomes for initial external arginine concentrations ranging from 1 to 100 μM and initial internal ornithine concentrations of **a**, 10 mM; **b**, 5 mM; **c**, 2.5 mM; and **d**, 0.5 mM.

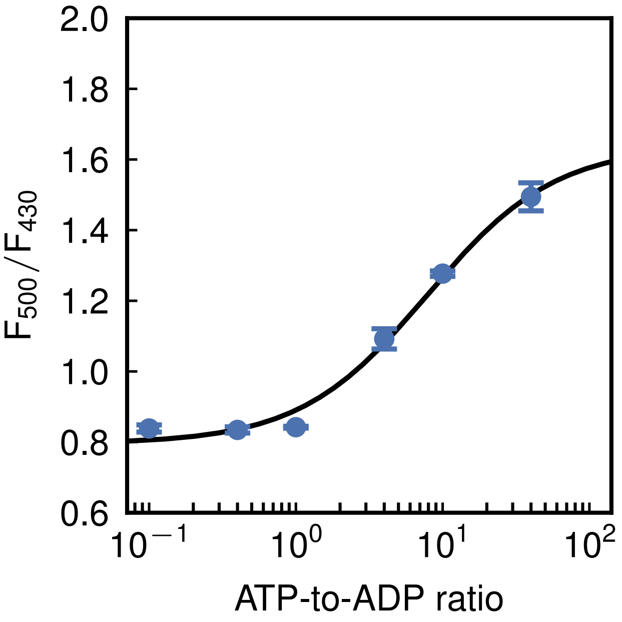


**Supplementary Figure 9: Calibration curve of PercevalHR in vesicles.** The calibration curve was obtained from the $F_{500}/F_{430}$ excitation ratio (500 and 430 nm) measured at defined ATP-to-ADP ratios, with total [ADP] + [ATP] maintained at 10 mM. Data were fitted with a Hill equation with Hill coefficient fixed to 1 (black line), as described in the Methods (start = 0.795, end = 1.638, $k$ = 7.9).

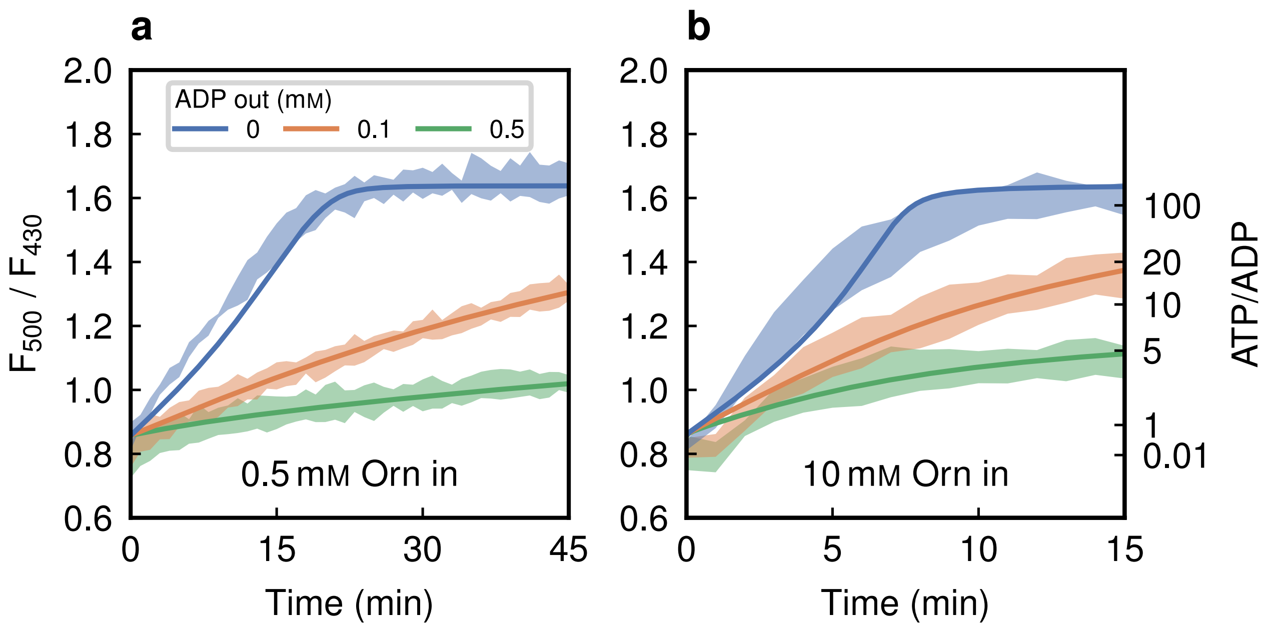


**Supplementary Figure 10: ADI pathway model reproduces experimental data with good agreement.** PercevalHR ratiometric signal ($F_{500}/F_{430}$) for the ADI pathway reconstituted in synthetic vesicles (same data as in Fig. 3). Solid lines show the model obtained from a single fit to the data.

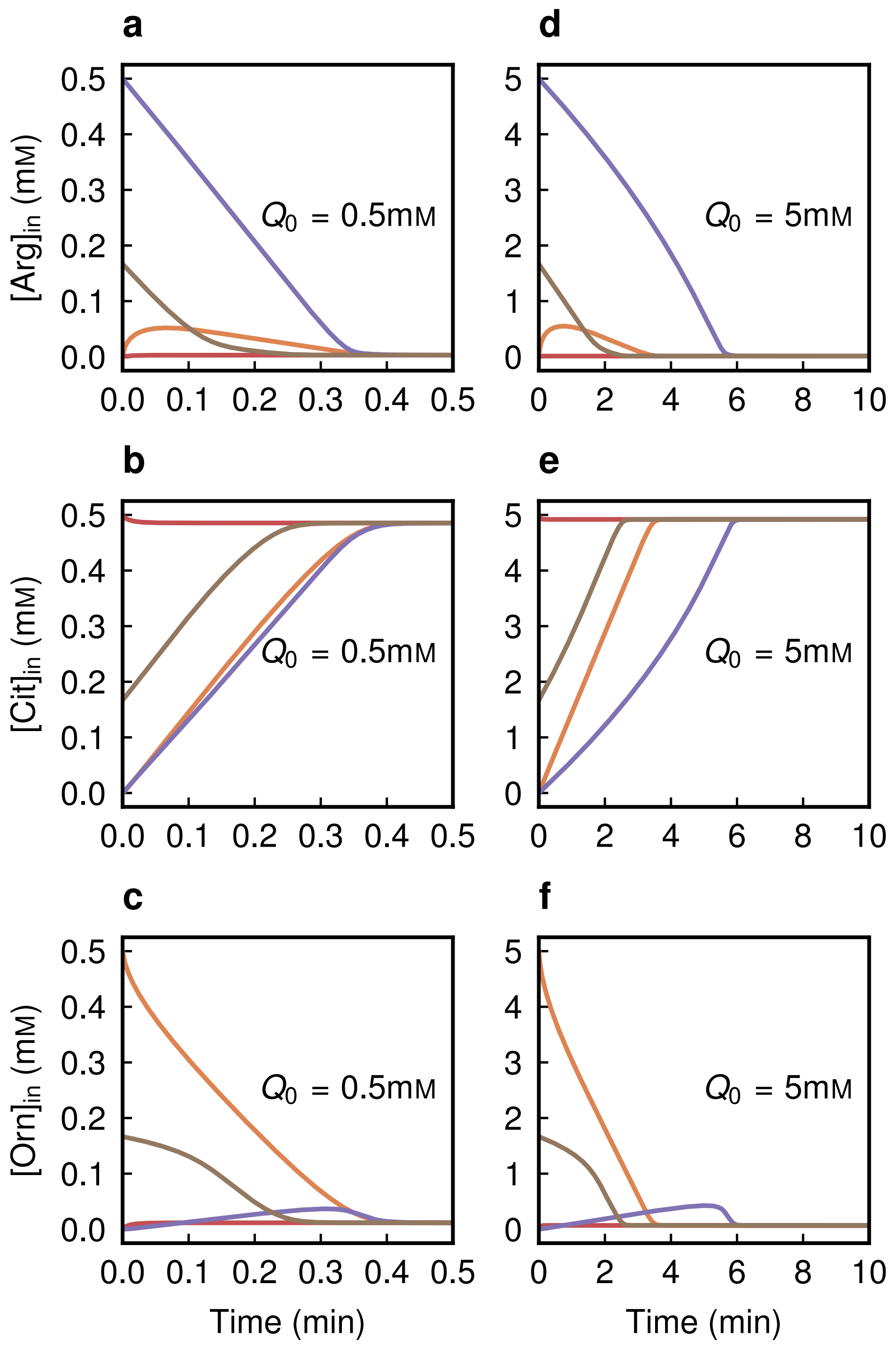


**Supplementary Figure 11: Transient metabolite dynamics depend on initial conditions but converge to a $Q_0$-determined steady state. a–c**, Time evolution of intravesicular arginine (**a**), citrulline (**b**), and ornithine (**c**) concentrations at $Q_0$ = 0.5 mM, for four initial conditions: all arginine ($[\mathrm{Arg}]^0_{\mathrm{in}} = Q_0$), all citrulline ($[\mathrm{Cit}]^0_{\mathrm{in}} = Q_0$), all ornithine ($[\mathrm{Orn}]^0_{\mathrm{in}} = Q_0$), and an equal split ($[\mathrm{Arg}]^0_{\mathrm{in}} = [\mathrm{Cit}]^0_{\mathrm{in}} = [\mathrm{Arg}]^0_{\mathrm{in}} = Q_0/3$). **d–f**, Same as **a–c** but at $Q_0$ = 10 mM. In all simulations, external arginine and ADP concentrations were held fixed at 100 mM. To allow the system to reach a steady state, all external concentration time derivatives were set to zero, and $NH_3$ and $CO_2$ membrane diffusion, carbamate hydrolysis, and cyanate hydrolysis were assumed to instantaneous.

### Supplementary tables

**Supplementary Table 1:** Kinetic parameters resulting from the characterization of the antiporter and the three cytosolic enzymes that constitute the ADI pathway.

| Characterization | Rate law | Kinetic parameter | | | |
|---|---|---|---|---|---|
| | | Symbol | Value | Uncertainty | Unit |
| *L. lactis* ArcA 194 kDa Tetramer | Michaelis-Menten with uncompetitive inhibition | $k_{A}$ | 6.93 | 0.17 | $s^{-1}$ |
| | | $K_{Arg}^{mA}$ | 5.1 | 2.2 | µM |
| | | $K_{Arg}^{iA}$ | 3.2[a] | | mM |
| *L. lactis* ArcB 245 kDa Hexamer | Ordered bi-bi | $k_{B}^{fv}$ | 62 | 13 | $s^{-1}$ |
| | | $k_{B}^{ufv}$ | 89/6[b] | | $s^{-1}$ |
| | | $K_{CP}^{mB}$ | 0.898 | 0.044 | mM |
| | | $K_{CP}^{iB}$ | 1.37 | 0.21 | µM |
| | | $K_{Orn}^{mB}$ | 1.095 | 0.044 | mM |
| | | $K_{Cit}^{mB}$ | 0.255 | 0.036 | mM |
| | | $K_{Cit}^{iB}$ | 3.168 | 0.051 | mM |
| | | $K_{P_i}^{iB}$ | 109.00 | 0.47 | mM |
| | | $K_{B}^{eq}$ | 13.00 | 0.27 | $\times 10^{-6}$ |
| *L. lactis* ArcC1 73 kDa Dimer | Reversible Hill equation | $k_{C}^{fv}$ | 914 | 20 | $s^{-1}$ |
| | | $K_{CP}^{mC}$ | 0.618 | 0.059 | mM |
| | | $K_{MgADP}^{mC}$ | 2.54 | 0.17 | mM |
| *L. sakei* ArcD 53.7 kDa Monomer | Ping-pong | $k_{D}^{+}$ | 6.4 | 1.4 | $s^{-1}$ |
| | | $k_{D}^{-}$ | 87 | 14 | $s^{-1}$ |
| | | $K_{Arg}^{mD}$ | 28.7 | 3.3 | µM |
| | | $K_{Orn}^{mD}$ | 1.402 | 0.094 | mM |

[a] Pols et al.[3]
[b] Marshall and Cohen[4]

**Supplementary Table 2:** Spontaneous processes included in the model of the arginine breakdown pathway. The chemical constant serves as the parameter determining the kinetics of each process in the model.

| Reaction equation | Chemical constant | | | |
|---|---|---|---|---|
| | Symbol | Value | Unit | Ref. |
| $MgADP \rightleftharpoons Mg^{2+} + ADP$ | $K^{d}_{MgADP}$ | 0.6 | mM | 5 |
| $MgATP \rightleftharpoons Mg^{2+} + ATP$ | $K^{d}_{MgATP}$ | 0.1 | mM | 5 |
| $H_2PO_4^- \rightleftharpoons P_i + H^+$ | $pK^{a}_{buff}$ | 6.87[a] | | |
| $NH_4^+ \rightleftharpoons NH_3 + H^+$ | $pK^{a}_{NH_4^+}$ | 9.084[b] | | 6 |
| $CO_2 + H_2O \rightleftharpoons HCO_3^- + H^+$ | $pK^{a}_{CO_2}$ | 6.33 | | 7 |
| $CP \longrightarrow P_i + \text{Cyanate} + H^+$<br>$CP + H_2O \longrightarrow P_i + \text{Carbamate} + H^+$ | $k^{hyd}_{CP}$ | $1.5 \cdot 10^{-4}$ | $s^{-1}$ | This study |
| $\text{Carbamate} + H_2O \rightleftharpoons NH_3 + HCO_3^-$ | $k^{+}_{Carb}$ | 124[b] | $s^{-1}$ | 6 |
| | $K^{eq}_{Carb}$ | 0.53 | M | 8 |
| $\text{Cyanate} + 2\,H_2O \longrightarrow NH_3 + HCO_3^-$ | $k^{hyd}_{OCN^-}$ | $5.341 \cdot 10^{-4}$ | $s^{-1}$ | 9 |

[a] Calculated at an ionic strength of 0.1 M
[b] Calculated at $T = 30\,^{\circ}C$ using linear interpolation

**Supplementary Table 3:** Kinetic parameters resulting from fitting the ADI pathway model to experimentally measured $F_{500}/F_{430}$ ratios.

| Characterization | Rate law | Kinetic parameter | | | |
|---|---|---|---|---|---|
| | | Symbol | Value | Uncertainty | Unit |
| *L. lactis* ArcC1 73 kDa | Reversible Hill equation | $k_{C}^{ufv}$ | 573 | 40 | $s^{-1}$ |
| | | $K_{Carb}^{mC}$ | 44 | 10 | µM |
| | | $K_{MgATP}^{mC}$ | 9.5 | 1.7 | mM |
| Mitochondrial AAC 33 kDa | Ping-pong | $k_{T}^{+}$ | 2.45 | 0.14 | $s^{-1}$ |
| | | $k_{T}^{-}$ | 8.5 | 1.7 | $s^{-1}$ |
| | | $K_{ADP}^{mT}$ | 2.83 | 0.13 | µM |
| | | $K_{ATP}^{mT}$ | 4.05 | 0.91 | µM |
